# Disorganization of the histone core promotes organization of heterochromatin into phase-separated droplets

**DOI:** 10.1101/473132

**Authors:** S. Sanulli, MJ. Trnka, V. Dharmarajan, RW. Tibble, BD. Pascal, A. Burlingame, PR. Griffin, JD. Gross, GJ. Narlikar

## Abstract

The heterochromatin protein HP1 is proposed to enable chromatin compaction via liquid droplet formation. Yet, a connection between phase separation and chromatin compaction has not been experimentally demonstrated. More fundamentally, how HP1 action at the level of a single nucleosome drives chromatin compaction remains poorly understood. Here we directly demonstrate that the S. *pombe* HP1 protein, Swi6, compacts arrays of multiple nucleosomes into phase-separated droplets. Using hydrogen-deuterium exchange, NMR, and mass-spectrometry, we further find that Swi6 substantially increases the accessibility and dynamics of buried histone residues within a mononucleosome. Restraining these dynamics via site-specific disulfide bonds impairs the compaction of nucleosome arrays into phase-separated droplets. Our results indicate that chromatin compaction and phase separation can be highly coupled processes. Further, we find that such coupling is promoted by a counter-intuitive function of Swi6, namely disorganization of the octamer core. Phase separation is canonically mediated by weak and dynamic multivalent interactions. We propose that dynamic exposure of buried histone residues increases opportunities for multivalent interactions between nucleosomes, thereby coupling chromatin compaction to phase separation. We anticipate that this new model for chromatin organization may more generally explain the formation of highly compacted chromatin assemblies beyond heterochromatin.

## Main text

Heterochromatin formation impacts genome function at multiple scales. Thus heterochromatin enables heritable gene repression, helps maintain chromosome integrity and provides mechanical rigidity to the nucleus^1,2^. It has been proposed that these diverse functions arise in part from compaction of the underlying chromatin. Yet how heterochromatin proteins enable chromatin compaction remains poorly understood.

A major type of heterochromatin contains at its core the complex formed between HP1 proteins and chromatin that is methylated on histone H3, lysine 9 (H3K9me)^3^. Previous work has suggested that dimerization and higher-order oligomerization of HP1 proteins helps promote chromatin compaction^4,5^. More recently it has been shown that human and *Drosophila* HP1 proteins can form phase-separated droplets and that chromatin can get incorporated into droplets formed by the human HP1 protein HP1α^6,7^. These results have led to the model that phase separation mediated by HP1 proteins compartmentalizes and compacts chromatin.

Further, it has long been theorized that macromolecular crowding within the nucleus may lead to phase separation of chromatin^8^. However, two types of fundamental questions have remained unanswered: (1) It is poorly understood how HP1 proteins regulate chromatin structure to enable its compaction and, (2) it is not clear whether and how chromatin compaction mechanisms are related to phase separation. Importantly, experimental evidence indicating that HP1 mediated phase separation enables chromatin compaction has been lacking.

Here we address the above questions by assessing the impact of the S. *pombe* HP1 protein, Swi6, on nucleosome conformation and by investigating the relationship between chromatin compaction and phase separation. We used a combination of complementary biophysical methods to study how Swi6 binding affects dynamics at the atomic level within a mononucleosome. Because HP1 proteins are proposed to mediate gene silencing in heterochromatin by lowering chromatin accessibility, we expected that internal motions within a nucleosome would be reduced upon Swi6 assembly^9^. Instead, we find that Swi6 binding results in increased dynamics and increased solvent accessibility of buried histone residues. We further find that Swi6 can promote self-association of nucleosome arrays and such self association driven compaction manifests as droplet formation. Intriguingly, the increased accessibility and dynamics of the histone core enabled by Swi6, promotes chromatin compaction into liquid droplets. Overall an unexpected model for heterochromatin assembly arises from this work wherein HP1 proteins act as nucleosome disorganizing agents so that compaction of chromatin is tightly coupled to its phase separation.

### Swi6 binding to octamer surface increases dynamics and accessibility of buried regions

Swi6 has two structured domains, the chromodomain (CD), which binds the H3K9me mark and the chromoshadowdomain (CSD), which forms a dimer that can bind certain types of hydrophobic protein sequences (Fig.1a)^10^. Additionally, Swi6 has an unstructured hinge region connecting the CD and CSD. The hinge is enriched for lysines and proposed to bind DNA in a sequence non-specific manner^11^. Previous work has indicated that four molecules of Swi6 bind to a single H3K9me nucleosome and that Swi6 makes contacts with the nucleosome core in addition to contacts made between its CD and the H3K9methylated histone tail^12,13^. To better understand the nature of these additional contacts, we performed cross-linking mass spectrometry (XLMS) on the Swi6-nucleosome complex using nucleosomes containing a methyl lysine analog on H3K9 (H3Kc9me3 nucleosomes). We used the zero-length reagent EDC, which cross-links the carboxylate side chains of Asp and Glu to amino side chains of Lys. Numerous cross-links were observed between histones and conserved Swi6 domains including the CD, CSD, and the hinge (Fig. 1b, c, Extended Data Figs 1a, b).

**Fig. 1:**
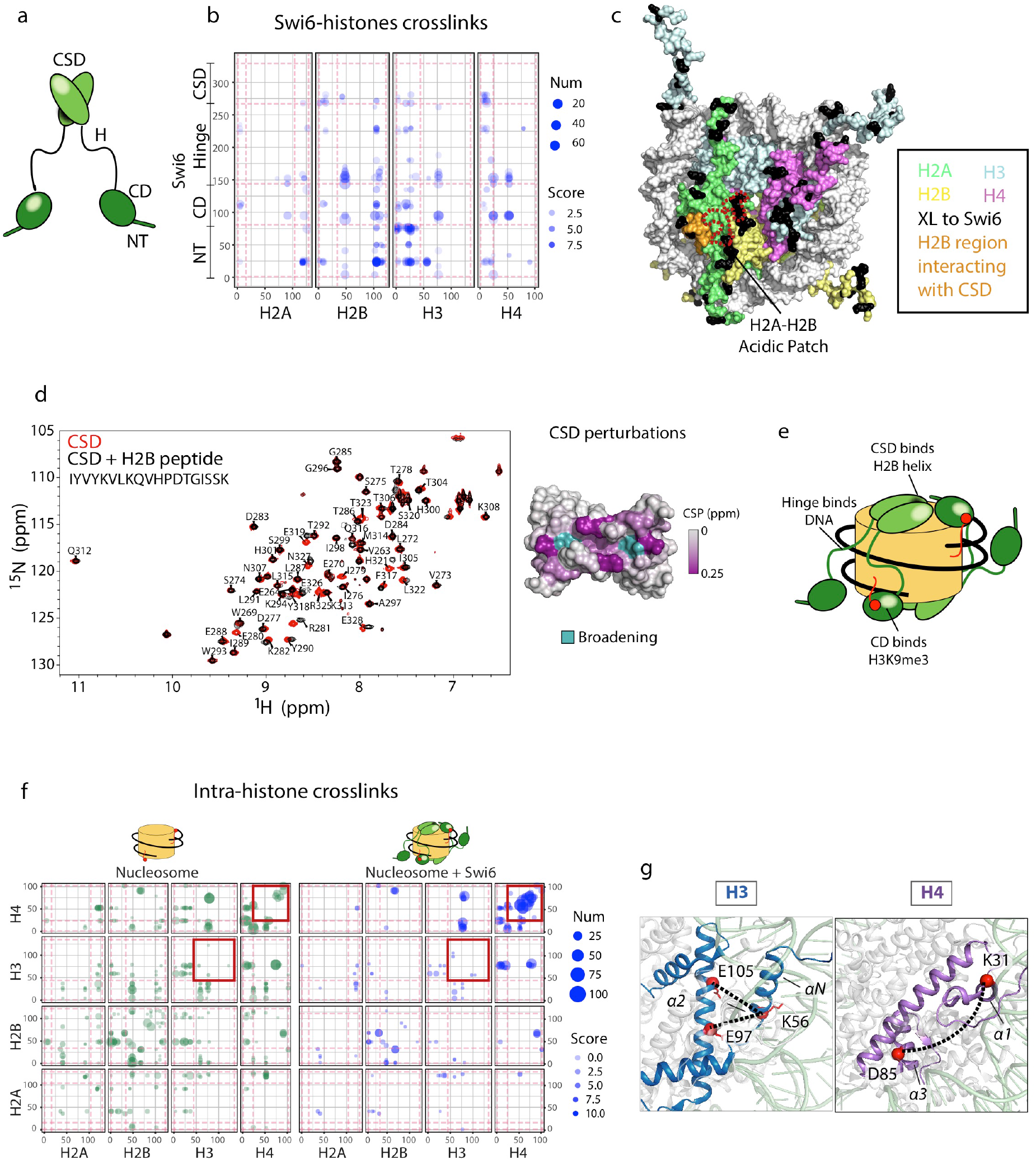
Swi6 makes several contacts with the histone octamer core and alters the pattern of intra-histone cross-links. (a) Domain architecture of Swi6. The chromodomain (CD), chromoshadowdomain (CSD), N-terminal region (NT) and hinge (H) are shown. (b) Analysis of EDC cross-links between histones and Swi6 domains. Cross-links are plotted as filled blue circles with area proportional to the number of MS2 spectra identifying a given cross-link. The numbers indicate the residue positions. N- and C-terminal histone tails are demarcated from the core regions by the dashed pink lines. The different Swi6 domains are also analogously demarcated. The H3 tail cross-links to the NT and CD of Swi6; the C-terminus of H2A cross-links to Swi6 NT. Several cross-links between Swi6 and histone H4, especially in its tail, are detected. (c) The histone residues that cross-link to Swi6 are mapped in black on the nucleosome structure. The different histones are colored as indicated. The H2A-H2B acidic patch is indicated with red dashed lines. The H2B regions interacting with the CSD is colored in orange. (d) The Swi6 CSD binds to H2B core. Left panel: superimposition of 1H-15N HSQC spectra of Swi6 CSD with (black) and without (red) H2B peptide. The sequence of the H2B peptide is shown in black. Right panel: the Swi6 CSD crystal structure colored by chemical shift perturbation upon the addition of 2X molar ratio of H2B peptide. In teal are the residues that show resonance broadening beyond detection upon peptide addition (PDB 1E0B). (e) Model for how Swi6 engages the nucleosome. (f) Cross-linking between histone proteins is altered by Swi6 binding. Left panel: Analysis of intra-histone cross-links. Each axis maps residue position within a histone, while the red dashed lines indicate the boundaries between histone tail and core domains. A cross-link is represented by a filled circle between two residues, with area proportional to the number of MS2 spectra identifying a given cross-link. Cross-links found in the free nucleosome state are in green, while cross-links found in the Swi6 bound nucleosome are in blue. The red squares highlight the changes in the H3-H3 and H4-H4 cross-linking patterns between the free and bound nucleosome states. (g) Intra-histone cross-link changes due to Swi6 binding are shown on the nucleosome structure. Histone H3 and H4 are colored in blue and purple, respectively. Examples of residues found cross-linked only when Swi6 is present are represented as red spheres.

The highest density of cross-links between the nucleosome and Swi6 arises from histone H3, followed by H2B (core), H4 (core and N-tail), and H2A (C-terminus). The H3 cross-links are predominantly between the H3K9me3 tail and N-terminus of Swi6, as anticipated by prior studies (Extended Data Fig. 1b)^14,15^. The cross-linking patterns show that Swi6 interacts extensively with the disk face of the octamer core, particularly through H2B. Histone H2B, together with histone H2A, forms the nucleosome acidic patch on the disk face, which is involved in binding several chromatin regulators^16^. While we observed some cross-links to acidic patch residues, the majority of the acidic patch residues did not cross-link to Swi6, suggesting that this frequently used “hotspot” for nucleosome interactors remains accessible even after Swi6 binding (Fig. 1c).

The CSD-CSD dimer interface of Swi6 has previously been implicated in binding the nucleosome core, but the region on the nucleosome that is contacted by the Swi6 CSD was not known^12,17^. Our cross-linking data suggests that the CSD interacts with the octamer core through histone H2B. Specifically, residue K43 in H2B cross-links with residues D278 and E281, which are located near the CSD dimerization interface (Fig. 1c). This interface is known to interact with proteins through short linear sequences containing the motif ϕx(V/P)xϕ (where ϕ and x, indicate a hydrophobic and any amino acid, respectively)^18–20^. The amino acid sequence surrounding K43 is rich in hydrophobic amino acids and is located on the surface of the nucleosome. We reasoned that this region of H2B, which lies at the C-terminal of the α1 helix, could bind the CSD dimer cleft as a short linear motif. To test this hypothesis, we performed ^1^H-^15^N HSQC NMR with ^15^N-labeled CSD in the presence of an unlabeled H2B peptide that corresponds to residues 36 to 54. Binding of the H2B peptide causes chemical shift perturbations (CSPs) in the CSD cleft (Fig. 1d, Extended Data Fig. 1c). The CSPs map on to the region of the CSD that has been shown to interact with specific peptides from the Shugoshin and Clr3 proteins, which are known ligands of Swi6^19^. Interestingly, in the available structures of the CSD in complex with peptides, the ϕx(V/P)xϕ motifs adopt a linear unfolded conformation to fit into the pocket of the CSD dimer^20,21^. It is therefore likely that, in order to bind the CSD, at least a portion of the H2B α1 helix undergoes a structural transition to become a short linear motif. Together, these experiments demonstrate that Swi6 establishes extensive contacts with the nucleosome histone core in addition to the H3K9 methylated tail (Fig. 1e).

While identifying Swi6-histone cross-links we also compared the intra-nucleosomal histone-histone cross-links observed in the nucleosome alone to those observed in the Swi6-bound state. Many of the intra-nucleosomal cross-links between core histone domains remained unchanged upon Swi6 binding. However several unique cross-links were detected in the Swi6-bound state between H3-H3 and H4-H4 (Fig. 1f). For instance, the buried residues E97 and E105 of histone H3, whose Cα residues are ~15 Å from K56, cross-link with K56 only in the presence of Swi6 (Fig. 1g). A large increase in intra-H4 cross-links is also detected in presence of Swi6, involving mainly residues in helices a2 and a3. Among them, D85 is a buried residue of H4 that cross-links with residue K31 of H4 (~30 Å away in terms of Ca-Ca distance) only with Swi6 bound (Fig. 1g, Extended Data Fig. 2a). These new intra-histone cross-links are unexpected as these residues are not within the standard distance captured by EDC cross-linking. Together with the possibility that CSD binding partially unfolds the H2B α1 helix (Fig. 1c, d), the presence of new intra-histone cross-links suggests that Swi6 binding perturbs the canonical conformation of the histone octamer. It is formally possible that the new cross-links capture transient inter-nucleosomal interactions mediated by Swi6. However, because the regions showing new cross-links in the Swi6-bound state are in buried regions of the nucleosome, any inter-nucleosomal cross-links would still require a substantial change in the solvent accessibility of these residues (Extended Data Fig. 2a).

The results above raised the possibility that Swi6 binding alters the conformational states accessible to the histone octamer, which could manifest as an alteration in histone conformational dynamics. To more directly test for such a possibility, we used hydrogen deuterium exchange with mass spectrometry (HDX-MS). This method measures the exchange of backbone amide hydrogens in the protein with solvent deuterium, which reports on protein backbone conformation, dynamics and solvent accessibility. Thus, a decrease or increase in deuterium uptake can often be interpreted respectively, as a decrease or increase in solvent exposure. HDX-MS has been used previously to assess the conformation of centromeric nucleosomes containing the H3 variant CENP-A^22^. In these studies, binding of the regulator protein CENP-C to CENP-A nucleosomes was found to decrease the deuterium uptake of internal histone residues, implying a rigidification of the histone core. Deuterium exchange was carried out as a function of time on H3Kc9me3 mononucleosomes alone or in complex with saturating concentrations of Swi6 (Extended Data Fig. 3a, see Methods). Under HDX conditions, peptic digest of nucleosomes yielded ~70% sequence coverage of the histone core and minimal coverage on the histone tails. Availability of overlapping peptic peptides allowed us to unambiguously assign the HDX-MS data to the histone core domains^23^.

Because HP1 proteins are proposed to decrease chromatin access, our simplest expectation was that Swi6 binding to a mononucleosome would decrease solvent access to the histone core resulting in decreased deuterium uptake^9^. Instead, Swi6 binding to H3Kc9me3 nucleosomes resulted in a robust and widespread increase of deuterium incorporation throughout the histone octamer (Fig. 2a-c, Extended Data Figs 3b, c). Specifically, more than a 35% increase in deuterium incorporation compared to nucleosomes alone is observed in: (i) residues 74-90 of H3, corresponding to a portion of helix a1 and a2, and the connecting loop 1; (ii) residues 48-61 of H3, corresponding to helix αN of H3; (iii) and residues 85-102 of H4, corresponding to helix α3. Other regions with a significant increase in deuterium uptake are the C-terminal region and helix α2 of H2A (residues 113-129 and 40-51), the C-terminal of H2B (helix αC, residues 104-122), and H4 helix α2 and loop 2 (residues 50-60 and 72-84) (Fig. 2c). These increases in the rates of deuterium incorporation caused by Swi6 binding indicate extensive changes in the backbone hydrogen bonding of histones within the nucleosome and partial unfolding of the helixes in the histones. Remarkably, the regions of histones H3 and H4 showing the highest degree of deuterium incorporation upon Swi6 binding are buried within histone-histone and histone-DNA interfaces in the canonical nucleosome structure (Figs 2d, e). These results indicate that (i) Swi6 binding increases the conformational states accessible to the histone octamer and (ii) the nature of the conformational changes involves a widespread increase in the solvent accessibility of buried histone residues.

**Fig. 2:**
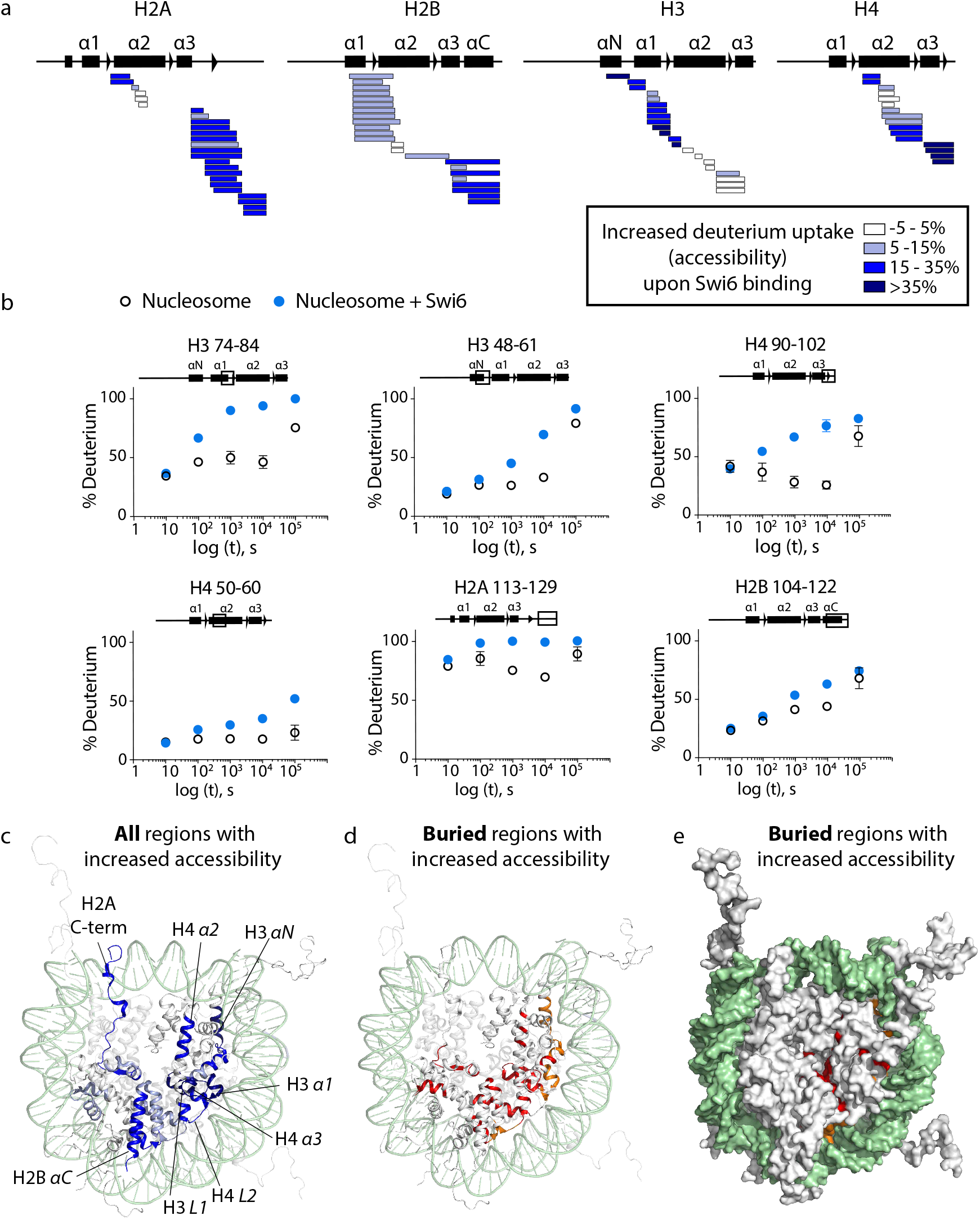
Swi6 binding causes an increase in solvent accessibility of buried histone residues. (a) Differential HDX-MS of all histone subunits of the nucleosome Swi6-bound versus free state from a single time point (10^4^ sec; see all time points in Extended Data Fig. 3). Each horizontal bar represents an individual peptide, and peptides are placed beneath a schematic depiction of secondary structure elements. The color map represents the percent increase in deuterium uptake observed in presence of Swi6, as indicated in the legend. (b) Kinetics of deuterium uptake of representative histone peptides over the time course. Data are the mean and SD of triplicates. For some points, the error bars are not shown as they are shorter than the height of the symbol. (c) Histones in the nucleosome structure (1KX5) are colored according to the % of deprotection shown in panel A. For clarity, only one copy of histones has been colored in the structure. DNA is shown in light green. (d) Buried histone regions that show an increase in deuterium uptake upon Swi6 binding are highlighted in ribbon diagram, with orange residues proximal to DNA and red residues proximal to histones. (e) Same residues in (d) shown in space-fill to visualize buried nature.

To further characterize the histone conformational dynamics we turned to methyl-TROSY NMR spectroscopy^24^. This approach allows (i) residue level resolution as opposed to peptide level resolution; (ii) investigation of side-chains as opposed to the protein backbone; (iii) monitoring changes in magnetic environment, due to binding or a conformational change; and (iv) determination of the time-scale of the side-chain dynamics. Specifically, this method monitors the chemical shift of methyl groups in macromolecular assemblies as large as the 26S proteasome and the nucleosome bound to the chromatin remodeler SNF2h^25–28^. Histones were labeled with ^13^C on methyl groups of isoleucine, leucine and valine (ILV) in an otherwise deuterated environment. These methyl groups are predominantly distributed in the hydrophobic buried core of the nucleosome (Fig. 3)^29^. To optimize coverage and spectral resolution, nucleosomes were labeled with isoleucine on histone H3 whereas histone H2B was labeled with leucine and valine. This labeling scheme maximizes the distribution of methyl probes across the nucleosome core and minimizes dephasing due to inter methyl-group dipolar interactions^30^.

**Fig. 3:**
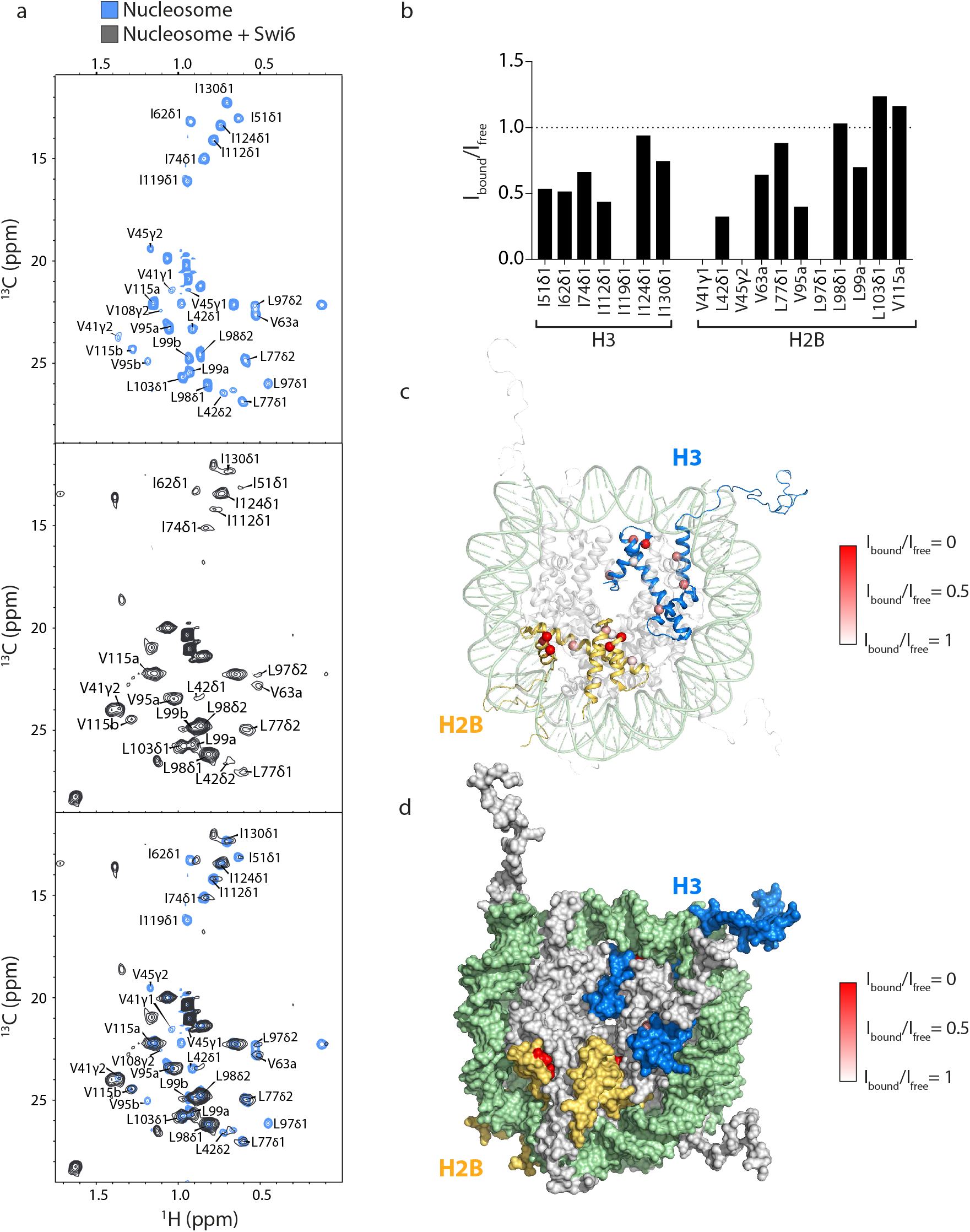
Swi6 binding increases protein dynamics within the octamer core. (a) The methyl-TROSY spectra of Ile-methyl-labeled H3 and Leu/Val-methyl-labeled H2B in the nucleosome alone (blue), and bound to Swi6 (grey). Three resonances in H2B are not assigned as they correspond to residues V15, V38 and V66 present only in the Xenopus histone. Some resonances are largely unaffected (I124, L98, L99, V115, and two not assigned), while others are broadened and a few resonances appear in presence of Swi6. (b) Quantification of cross-peak broadening for H3 and H2B residues. Top x-axis represents relative peak volumes of residues in the nucleosome-Swi6 complex (I_bound_) compared to those in the nucleosome alone (I_free_). The two sets of spectra were normalized using the volumes of their respective L98 and two non-assigned peaks, which do not show broadening upon Swi6 binding. To be conservative, only the unambiguously assigned peaks have been included in the quantification. Cross-peaks that disappear are taken to have an I_bound_ value of zero. (c) Ca of Ile residues in H3 and Leu/Val residues in H2B are shown as red spheres in the nucleosome structure. The intensity of red color represents the extent of broadening determined by I_bound_ /I_free_. (d) Surface representation of nucleosome in (c) indicating that most of the perturbed residues are buried.

The methyl-TROSY spectra of H3 and H2B are well resolved and small (<0.1ppm) changes in chemical shift are observed between the cross-peaks of our spectra and of the published spectra of *Drosophila* nucleosomes (Fig. 3a, Extended Data Fig. 4a)^28^. Due to the small changes and the high protein conservation between the two species, we were able to transfer the assignments to all the cross-peaks of H3 and most of the cross-peaks of H2B (Extended data Fig. 4b). Binding of Swi6 to the nucleosome induced shifts and broadening of select resonances (Fig. 3a, b). Because (i) Swi6 is deuterated, (ii) its binding affinity for the nucleosome is tight (K_d_ <100 nM), (iii) and nearly all of the ILV residues that undergo resonance broadening are buried in the canonical nucleosome structure, we interpret differential resonance broadening as intramolecular conformational dynamics on the ms-μs timescale in the histone octamer core induced by Swi6 (Figs 3c, d, Extended Data Fig. 5)^31^. Importantly, the changes in side chain dynamics detected by NMR are consistent with the changes in backbone solvent accessibility detected by HDX-MS and the changes in intra-histone cross-links (Extended Data Figs 6a, b). For example, H3 residues I51 and I62, which are among the most affected by Swi6 according to the methyl-TROSY NMR, are within the regions that show higher deuterium uptake. The NMR results also rule out the possibility that the increased solvent accessibility observed in the HDX-MS data is due to disassembly of the nucleosome. This is because the cross-peaks of free histones fall in a substantially different chemical shift regime than what we observe in the presence of Swi6.

It is informative to compare our observations here with previous methyl-TROSY NMR studies carried out with linker histones^28,32^. These studies showed that binding of a linker histone, which enables chromatin compaction, does not cause the types of ms-μs dynamics within the histone core as we observe with Swi6. We therefore propose that not all proteins that enable chromatin compaction will have the same effect on the nucleosome core.

In principle structural studies have the potential to inform upon the degree of deformation in the histone core. For example, to date there are two cryo-EM structures, one with Swi6 bound to a mononucleosome and one with human HP1α bound to a dinucleosome^12,33^. While the Swi6-nucleosome structure shows density that is compatible with four Swi6 molecules bound to a mononucleosome, the HP1α-dinucleosome structure shows density that is compatible with one HP1α dimer bridging across two nucleosomes. However, the resolution in both structures is not sufficiently high to detect the types of local histone dynamics implied by our results. We speculate that dynamics of nucleosome induced by heterochromatin proteins may limit the ability to obtain high-resolution structures, which remains a challenge for the future.

In summary, overlaying the results from the three complementary methods, XLMS, HDX-MS and methyl-TROSY NMR, shows a high degree of overlap in the histone regions that are perturbed upon Swi6 binding (Extended Data Figs 6a, b). Further, the XLMS and ^15^N NMR data indicate that in addition to the H3K9me3 tail, Swi6 interacts with the nucleosome octamer core, mainly through histone H2B. Moreover, at least a portion of the nucleosome acidic patch is available for interactions with additional binding partners. Most surprisingly however binding by Swi6 on the nucleosome surface increases the solvent accessibility and dynamics of residues within the nucleosome interior.

### Octamer dynamics promote recognition of the H3K9 methyl mark by Swi6

To test if the conformational changes in the octamer detected by HDX-MS, NMR and XLMS play a functional role in the assembly of Swi6 on nucleosomes, we used disulfide linkages that lock the octamer into its ground state conformation^27^. If conformational rearrangements within the octamer core are energetically coupled to Swi6 binding, then constraining these rearrangements is predicted to reduce Swi6 binding.

To test the role of dynamics at the histone H3-H4 interface, we restrained the dynamics by inserting cysteines at positions I62 of H3 and A33 of H4. This region is buried in the octamer, but showed major perturbations by NMR, HDX-MS and XLMS (Fig. 4a, Extended Data Fig. 6a). We were able to obtain nucleosomes containing nearly 90% disulfide-linked histones (Extended Data Fig. 7a). To make these nucleosomes, we used H3K9me3 that was enzymatically methylated (Extended Data Fig. 8, Methods). Swi6 exhibited ~5 fold weaker affinity for the H3K9me3 methylated nucleosomes containing one disulfide bridge between H3 and H4 (referred to as H3•H4 S-S) compared to the same nucleosomes with reduced cysteines (H3•H4S-H) (Extended Data Fig. 7b, Table S1). Similar results were obtained with nucleosomes in which the dynamics of H4 α2 helix were restrained with three disulfide links (Extended Data Figs 7c-f). Remarkably, the disulfide cross-links did not affect the binding of Swi6 to un-methylated nucleosomes (Extended Data Figs 7g, h). Thus, Swi6 displays a greater specificity for H3K9me3 methylated nucleosomes when the H3-H4 dynamics are not restrained. Together, these results indicate that conformational plasticity within the histone octamer is important for Swi6 to bind the nucleosome core particle in a manner that allows recognition of the K9me3 mark on histone H3.

**Fig. 4:**
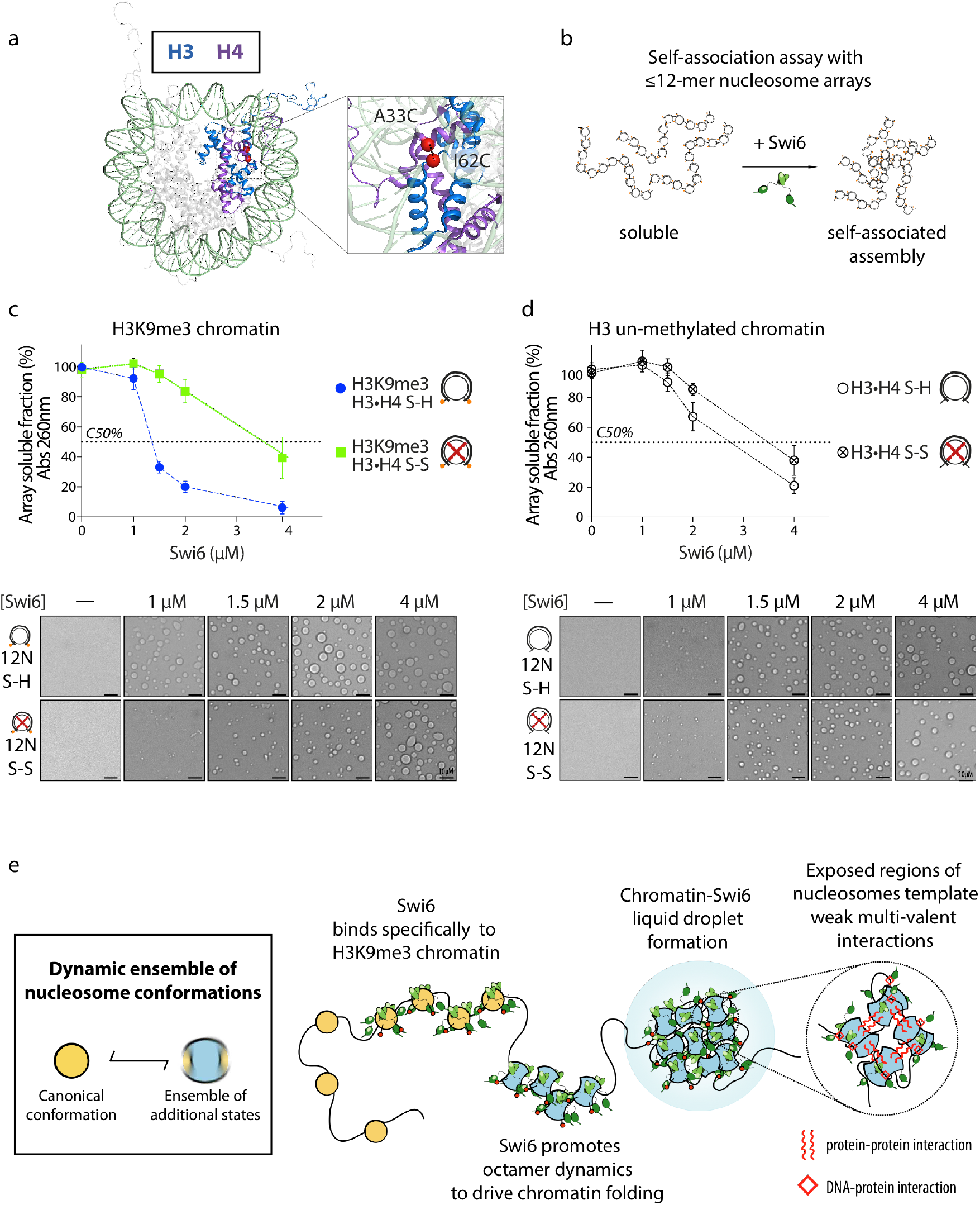
Nucleosome octamer dynamics promote Swi6-mediated chromatin condensation and phase separation. (a) Depiction of location of cysteine residues in H3 and H4 for generation of disulphide cross-links. (b) Schematic of chromatin self-association assay. (c) Precipitation assays performed on nucleosome arrays containing H3K9me3•H4 S-S (oxidized) and H3K9me3•H4 S-H (reduced) octamers as a function of increasing concentration of Swi6. The dotted line represents the C_50_, which is the concentration of Swi6 at which 50% of arrays are precipitated. Lower panel: bright field representative images of the array-Swi6 complex analyzed in the top panel, showing the formation of phase-separated droplets. (d) Nucleosome array precipitation assays as in (c), performed on un-methylated arrays containing H3•H4 S-S and H3•H4 S-H octamers. Lower panel: bright field representative images of the array-Swi6 complex analyzed in the top panel. (e) Model for Swi6-heterochromatin organization. Inset: nucleosomes in solution can sample an ensemble of conformational states, of which the canonical conformation observed in the crystal structure is the most populated and lowest energy state. Model: Swi6 exploits nucleosome plasticity to drive heterochromatin organization. Binding of Swi6 to H3K9me3 nucleosomes is coupled to loosening of histone-DNA and histone-histone contacts, which increasing the dynamics within the octamer core. This allows the nucleosome to more readily access a larger ensemble of dynamically interconverting conformational states. In these conformational states normally occluded regions of the octamer core become deprotected and available to establish transient weak multivalent interactions. Such interactions enable chromatin condensation into phase-separated liquid droplets. Red spheres represent the H3K9me3 mark; Swi6 dimers are shown in green.

### Increased octamer dynamics promote chromatin compaction and phase separation

All the results above describe effects of Swi6 in the context of a mononucleosome. We next asked how these effects at the level of mononucleosome impact the function of Swi6 in the context of compaction of nucleosome arrays. We used a classic *in vitro* method in which self-association is assayed by measuring the fraction of nucleosome arrays that precipitate in the presence of divalent metal ions or specific regulator proteins (Fig. 4b)^34^. Previous work has shown that human HP1 proteins promote the self-association of nucleosome arrays as measured by the chromatin precipitation assay^13,35,36^.

With the goal of assessing the role of histone core dynamics in chromatin compaction, we assembled 12 nucleosome-arrays containing one cysteine pair at the H3-H4 interface (H3I62C-H4A33C). We first measured array precipitation in presence of Swi6 under reducing conditions, when the dynamics at the H3-H4 interface are not restrained. We found that Swi6 causes precipitation of H3K9me3 nucleosome arrays in a dose dependent manner, analogous to previous observations with human HP1 proteins (Fig. 4c). To quantify the dose-dependence we estimated a value for C50, the concentration of Swi6 at which 50% of the chromatin is precipitated. Given the steepness of the Swi6 dose-dependence an exact C50 value was technically challenging to determine. The C_50_ value is lower for H3K9me3 arrays (between 1-1.5 μM), than for un-methylated arrays (between 2-4 μM), consistent with Swi6’s higher affinity for mono nucleosomes carrying the H3K9me3 mark (Fig. 4c, d, Extended Data Figs 7b, e, g, h). However we note that the shape of the concentration dependence for the methylated vs. unmethylated arrays is also different, suggesting that in the absence of the H3K9me mark, the effect of Swi6 is less cooperative.

Next, we investigated if Swi6 on its own displays phase separation behavior. On its own, Swi6 remains in a clear solution at concentrations up to 4 μM analogous to results with unmodified human HP1α (Extended Data Fig. 9a)^7^. However when we investigated the Swi6-array precipitates under a microscope, we observed the formation of liquid droplets in a Swi6-dependent manner that correlates with increased array precipitation (Figs. 4c, d, lower panels). The presence of histones in the droplets was confirmed using fluorescently labeled histones (Extended Data Fig. 9b). Notably, Swi6 can also form droplets in presence of DNA alone, similar to what was shown for HP1α (Extended Data Fig. 9c)^7^. However, compared to chromatin, a higher Swi6 concentration is required to observe droplet formation with naked DNA. Further, precipitation was not observed with Swi6-mononucleosome complexes at the substantially higher concentrations used for the NMR experiments (see Methods for more details). These results indicate that (i) Swi6 mediated chromatin self-association assays report on the formation of phase-separated droplets, (ii) chromatin self-association and droplet formation can be coupled processes.

We then assessed the effects of restricting histone octamer dynamics on Swi6-mediated chromatin self-association and droplet formation by forming the disulfide-link between H3I62C and H4A33C. The site-specific disulfide-linkage impairs Swi6’s ability to precipitate H3K9me3 nucleosome arrays and to form droplets (Fig. 4c, C_50_ =1-1.5 μM vs. ~4μM). In contrast, this disulfide bond does not substantially affect Swi6 mediated precipitation of un-methylated arrays (Fig. 4d). These results suggest that histone core dynamics are important for Swi6-mediated self-association of chromatin fibers in a manner that is specific to the H3K9me3 mark.

We note here that the quality of the arrays used in these studies had to be carefully controlled to avoid over assembly of histone octamers as we noticed that over-assembled arrays displayed aberrant and non-reproducible precipitation behavior (Extended Data Fig. 9d, see Methods for details).

There are two types of models that could explain the effects of the disulfide cross-link: (i) Swi6 binds more weakly to the cross-linked nucleosomes as observed in Extended Data Figure 7b, and therefore a higher concentration of Swi6 is required to form droplets; (ii) increased dynamics within the histone core directly impact chromatin compaction. In terms of a binding effect, even at concentrations of Swi6 (1.5 and 2 μM) that are saturating and above the K_d_ for the corresponding cross-linked mononucleosome (K_d_~ 0.3 μM, Table S1), we see less precipitation and smaller droplet formation for cross-linked nucleosomes in comparison to the uncross-linked nucleosomes. These data suggest that restricting octamer dynamics has effects beyond affecting Swi6 binding. However, to more directly interrogate the second possibility, we asked if intrinsic chromatin compaction also relies on octamer core dynamics. Several prior studies have used the effects of Mg^2+^ to infer the intrinsic self-association properties of nucleosome arrays^34^. We therefore asked if the Mg^2^+ driven self-association of chromatin was affected by constraining octamer dynamics. We found that Mg^2+^ driven self-association of chromatin was also impaired by the site-specific disulfide-linkage between H3 and H4 (Extended Data Fig. 9e). Further investigation of the Mg^2^+ driven chromatin precipitates revealed the presence of liquid droplets suggesting that intrinsic chromatin compaction can also be coupled to phase separation (Extended Data Fig. 9f).

Overall the results indicate that Swi6 binding to H3K9 methylated chromatin increases the solvent accessibility and dynamics of buried histone residues, which in turn promotes chromatin compaction and droplet formation.

### Conclusions and implications

Chromatin compaction is often considered a hallmark of heterochromatin. Yet at a biophysical level how the assembly of heterochromatin proteins such as HP1 enables chromatin compaction is poorly understood. Current models imply that HP1 molecules can promote chromatin compaction by bridging across nucleosomes either as dimers or oligomers^4,5,13,35,36^. At the same time, chromatin compaction is proposed to involve interactions made by histone tails with near-by nucleosomes^37^. Both types of chromatin compaction models treat the octamer core as a largely rigid and passive unit with the canonical conformation observed in crystal structures. Our work uncovers another fundamental regulator of chromatin compaction that is missing from both types of models, namely structural plasticity of the octamer core. As we discuss below, a role for octamer plasticity leads to new models for the function of HP1 proteins and new models for the higher-level organization of chromatin.

Our results indicate that Swi6 is more than just a bridging molecule that recognizes the H3K9me3 mark. We find that Swi6 establishes many contacts with the folded histone core, including between the CSD-CSD dimer and histone H2B, and promotes increased conformational dynamics within the histone octamer. The increased accessibility and dynamics of buried histone residues may appear paradoxical when viewed from the repressive role of heterochromatin. However, the paradox is resolvable through a model in which chromatin compaction and phase separation are coupled. In well-studied model systems the formation of phase-separated assemblies is shown to rely on weak multivalent interactions between the component macromolecules^38,39^. Building on these studies we propose that the intrinsic folding of chromatin into higher-level assemblies is promoted when buried histone core residues become transiently more accessible (Fig. 4e). Such transiently exposed regions can template weak and multivalent interactions between nucleosomes to drive chromatin compaction and droplet formation. Our results in Extended Data Figure 9e are consistent with this possibility as we find that inhibiting octamer dynamics impairs intrinsic chromatin compaction. Within the above framework, HP1 proteins such as Swi6 would then promote chromatin compaction by substantially increasing intrinsic histone core dynamics and accessibility. Specifically, we propose that through extensive contacts with the whole nucleosome, including the H3 tail, the H2B α1 helix and nucleosomal DNA, Swi6 loosens but does not disassemble the overall nucleosome allowing greater dynamics within the histone core (Fig. 1e). Thus, by the framework described above, even though Swi6 increases dynamics at the level of individual nucleosomes, the net effect at the level of multiple nucleosomes is their sequestration and compaction into phase-separated droplets. The intrinsic ability of Swi6 to form oligomers could contribute additional modes of multivalency, further stabilizing the phase^4,5,12^. Such a model is consistent with previous observations of lower histone turnover in S. *pombe* heterochromatin and lower DNA accessibility in *Drosophila* heterochromatin^9,40^.

Dynamics within nucleosomes and heterochromatin have been previously described. In the context of such previous work, our studies provide a complementary set of observations. In the context of nucleosomes, seminal prior studies have shown that nucleosomal DNA unwraps and rewraps on the order of milliseconds^41^. Here we show that the dynamic conformational changes that are possible within an intact nucleosome extend deep into the histone core to reveal new surfaces that we propose help drive higher-level chromatin folding. Further, recent cryo-EM studies of nucleosomes have identified conformations in which perturbations in the histone helices are observed in nucleosomes that have partially disrupted DNA^42,43^. It is therefore possible that the octamer core alterations we observed are coupled to changes in DNA dynamics such that stretches of exposed DNA also participate in the interactions that drive phase separation. In the context of heterochromatin, previous work has indicated that in cells, HP1 molecules display “on-off” dynamics on time scales ranging from milliseconds to seconds^44–46^. Correspondingly *in vitro*, HP1 molecules have been shown to transiently stabilize more compact chromatin states^4,47^. Here we describe intra-molecular dynamics within the histone octamer when HP1 is bound. It is possible that intra-octamer dynamics are coupled to the residency time of HP1 binding, further adding to the transient nature of the multivalent interactions within a phase-separated state.

Several prior studies have indicated the absence of repeating and well defined higher-level chromatin structures in cells^48–52^. Thus, while defined chromatin structures such as the 30 nm fiber are observed in certain terminally differentiated cells, these structures have not been detected in a majority of the cell types that have been investigated^48,53,54^. A phase separation based mechanism for chromatin compaction and compartmentalization can explain the absence of long-range order within compacted chromatin in cells. Additionally, the specific phase separation based model that we propose opens up new regulatory possibilities. For example, we find that deformation of the octamer core by Swi6 is energetically coupled to recognition of the H3K9me3 mark. Therefore, the presence of H3K9me marks can help tune the nature of inter-nucleosomal interactions made within the chromatin phases by tuning the extent of octamer deformation. Analogously, specific ligands of Swi6, which bind its CSD-CSD interface and compete for its interaction with the nucleosome, can also regulate Swi6-driven chromatin compaction and phase separation. It also is possible that in some cases histone modifications and histone variants regulate chromatin compaction through direct effects on histone octamer plasticity. Finally we speculate that other highly compacted chromatin states such as mitotically condensed chromatin may also rely on a disorganized and dynamic histone octamer to achieve a high degree of compaction. Future studies will uncover how broadly histone plasticity and phase separation mechanisms regulate chromatin states and genome organization.

## Methods

### Protein expression, purification and isotope labeling

All the histones were expressed and purified from E. coli following published protocols ^55^ with some modifications for the isotopically labeled histones. For expression of deuterated histones, M9 minimal media was made in deuterium oxide (D2O, Cambridge Isotope Ltd., CIL). The media contained 1,2,3,4,5,6,6-D7 Glucose (CIL or Sigma-Aldrich) as the sole carbon source. For labeling the histones at ILV residues, ^13^CH_3_-methyl group α-ketoisobutyrate and α-ketoisovalearate (CIL) were added to the M9 media as precursors ^56^. Deuteration and ILV labeling of the histones were confirmed by mass spectrometry. All the mutant versions of the histones were made by quick-change site-directed mutagenesis or the Gibson Assembly method. All mutations were confirmed by sequencing the plasmids.

Methyl lysine analogue (MLA) containing H3 histones at position 9 (H3Kc9me3) were prepared as described previously^57^.

N-terminally 6X-His tagged Swi6 was purified from an *E. coli* as previously described^5^. Briefly, Swi6 was affinity purified with cobalt beads followed by TEV protease treatment to cleave the N-terminal 6x-His tag. After TEV cleavage, it was subjected to HiTrap Q HP column (GE Healthcare) and Superdex 200HR 10/300 column (GE Healthcare). Swi6 is stored in 25 mM HEPES pH 7.5, 150 mM KCl, 1mM DTT and 10% glycerol. For expression of deuterated Swi6, M9 minimal media was made in deuterium oxide (D2O, Cambridge Isotope Ltd., CIL) with 1,2,3,4,5,6,6-D7 Glucose (CIL) as the only carbon source.

^15^N-labelled CSD was expressed in M9 minimal media containing ^15^N-ammonium chloride as the sole nitrogen source. The CSD was purified as previously described^19^.

GST-Dim5 plasmid was kindly provided by Dr. Isao Suetake. GST-Dim5 was expressed in Rosetta cells in presence of 10 μM ZnSO_4_ as previously described^58^. Cells were lysed in 20 mM Tris pH 9.6, 500 mM NaCl, 0.05% NP40, 1 mM DTT and protease inhibitor EDTA-free. The protein is first purified on a GSTrap FF column (GE Healthcare), then on HiTrap Q HP column. Purified Dim5 is stored in 20 mM Tris pH 9.6, 150 mM NaCl, 1mM DTT, 5% glycerol.

### H3K9 enzymatic methylation

Dim5 was dialyzed in the Methylation Buffer (MB): 100 mM Tris-Cl pH 9.6, 20 μM ZnSO_4_, 2 mM β-mercaptoethanol, 10 mM KCl, 10mM MgCl_2_) plus 80 mM NaCl. Histone H3 C110A, I62C was resuspended in MB buffer at 0.1 mg/ml. The reaction condition was adapted from Mishima et al.^17^ and carried out at room temperature for 3 hours with 1:2 molar ratio histone:Dim5 in presence of 50 μM S-adenosylmethionine (SAM, NEB). The reaction pH is adjusted to 9.8 after SAM addition. The reaction is then mixed with 4 volumes of 9M deionized urea, and add reagents to reach 200 mM NaCl, 20 mM NaAc pH 5.2, 1 mM EDTA, 5 mM β-mercaptoethanol as final concentration. The sample pH is adjusted to 5.2 before loading it on a HiTrap SP HP cation exchange chromatography column (GE Healthcare), and elute with a salt gradient. The histone is collected, dialyzed in H_2_O 5mM β-mercaptoethanol and diluted 10x with 0.1 % formic acid for Mass Spectrometry analysis.

Methylation was followed using a LC-MS-ETD-MS assay of the intact histone. The protein was separated over a 15 cm x 75 μm ID PepMap C18 column (Thermo) using a NanoAcquity UPLC system (Waters) running a gradient from 10-75% B (acetonitrile + 0.1% formic acid) at 300 nl/min over 40 minutes. Analysis was performed by an online Orbitrap Fusion Lumos (Thermo) mass spectrometer. Precursor ions were measured at 120,000 resolution in the Orbitrap (AGC: 1e6). Ions that were 15-22+ charged with intensity greater than 1e6 were isolated in the quadrupole (1.8 m/z isolation window) and fragmented by ETD for 4 msec. Product ions were measured in the Orbitrap at 30,000 resolution (AGC: 9e5, max injection time: 250 msec, 3 microscans).

Precursor ion spectra were deisotoped using Xtract (Thermo). The deconvoluted M+H of the untreated histone (15220.442) was consistent with the expected mass of the H3-I62C mutant (15220.554) to 7.4 ppm and was about 11% oxidized on Methionine. Enzymatic methylation produced a pattern of deconvoluted M+H signals spaced 14 Da apart, although some of these signals were off by ±1 Da due to challenging monoisotopic peak detection. These were confirmed as methylations by bottom-up proteomic analysis of Lys-C digested sample. The methylated sample had no detectable unmethylated, monomethylated, or dimethylated species. All of the ions signals for which the ETD spectra were available showed 100% trimethylation at H3K9 based on their c-ions. Additional methylations could be localized to H3K18 and K3K14 (Extended Data Fig. 8).

### Mononucleosome Assembly

Histone octamer was assembled from purified histones by salt dialysis as described1. The 601 positioning sequence (147 bp) DNA fragment was made using restriction enzyme digestion of a plasmid carrying multiple copies of 601 DNA fragment as described previously^55^. Fluorescence DNA was generated by PCR amplification using a primer 5’-labelled with 5,6 carboxy-fluorescein (IDT) followed by gel purification^14^. Nucleosomes were assembled using published gradient dialysis based protocols^55^. The purification of the nucleosomes was carried out on a 10 to 30% glycerol gradient. For NMR experiments, the glycerol gradient step was replaced by gel filtration purification with a Superdex-200 column.

Disulfide cross-linked nucleosomes were made using histone Cys variants, see Table S1 and S2. In order to maximize the likelihood of disulfide bridge formation, the cysteine pairs were selected so as to have an interatomic distance ≤5 Å between the sulfur atoms facing each other, to have side chains pointing toward each other in the structure and to form bonds in Coot (https://www2.mrc-lmb.cam.ac.uk/personal/pemsley/coot/). All the cysteine variant histone proteins were purified in the presence of excess dithiothreitol (DTT) during all the steps of protein purification. To purify the cysteine variants, only the gel filtration step was carried out, as previously described^27^. For reduced nucleosome, the assembly and subsequent purification were carried out in the presence of 3mM TCEP in all the buffers. For disulfide-linked nucleosome formation, the octamer was subjected to oxidizing conditions. First, the octamer sample was diluted to a final octamer concentration of 0.5 mg/ml to minimize the formation of any intermolecular cross-links. Then it was subjected to 3 consecutive dialysis in refolding buffer (pH 8.5) in absence of any reducing agent as previously described^27^. Finally the solution containing octamers was mixed 4:1 with a solution containing 250 mM Tris-Cl pH 9, 2 M NaCl, 2.5 mM oxidized glutathione, 2.5 mM reduced glutathione and incubated 12 h at room temperature as previously described^59^. The sample was then dialyzed in refolding buffer with no reducing agent, before proceeding to nucleosome assembly. The cross-linking efficiency was confirmed by running the sample on a 15% SDS gel.

### 12 nucleosome-array assembly

DNA was generated by restriction enzyme digestion of a plasmid containing 12 consecutive 601s spaced by 20 bp, followed by gel purification. The reconstitution of nucleosome arrays followed the protocols described with some modifications^60^. Histone octamers are combined in equimolar amount with 12-mer DNA (12 repeats of the 601 DNA sequence separated by 20-bp linkers). Final dialysis against 10 mM TEK (10 mM KCl, 10 mM Tris pH 7.5, 0.1 mM EDTA) was performed overnight. After assembly, arrays were usually used for experiments within 1–2 d after assembly. Since the 601 repeats in the 12-mer DNA sequence are separated by HpaI restriction enzyme sites, quality of assembly was assessed by HpaI digestion followed by native gel. Over-assembly was avoided by ensuring that >95% of the digested fragments migrated as mononucleosomes rather than slower migrating species. Avoiding over-assembly was essential as we noticed that small amounts of super-stoichiometric histones caused precipitation in the absence of Swi6 and Mg^2^+ and also gave non-reproducible results.

### Precipitation assay

Swi6-mediated array precipitation assays were performed at 40 nM nucleosome arrays in 10 mM Tris, 0.1mM EDTA, 75 mM KCl, pH 7.8. Briefly, 5 μl of 2X array sample (80 nM) were incubated with 6 μL of 2X Swi6 solution for 20 min at 22 °C. Samples were centrifuged at 10,000 × g for 10 min at 22 °C, and the supernatants were transferred to a fresh tube. 5 μl of the supernatant was read at 260 nm. 3 technical replicates were performed for each precipitation experiment. Mg^2^+ precipitation assays were performed in TE 0.1 (10mM Tris pH 7.8, 0.1mM EDTA).

### Microscopy

Microscopy of the droplets was done on a Leica Axiovert 200M microscope using a 10X air objective. Samples were imaged on a 384 well plate, glass bottom, coated with Peg-silane (Laysan Bio). Image analysis was done in ImageJ.

### Fluorescence polarization binding measurement

Nucleosome polarization assays were conducted in buffer containing 0.02% NP-40, 150 mM KCl, 20 mM HEPES pH 7.5, 1 mM DTT at 22 °C. Each anisotropy sample contained a final nucleosome concentration of 5 nM and the Swi6 concentration was varied. The reaction was incubated 30 min at 22 °C and fluorescence polarization was measured on an Analyst HT (Molecular Devices). Data points from three independent Swi6 dilution curves were averaged and standard errors calculated. The following binding model was used to derive K_d_:

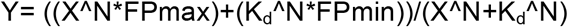

in which, Y is the fluorescence polarization signal observed; X is Swi6 concentration; FPmin is the fluorescence polarization signal for the probe alone; FPmax is the fluorescence polarization signal at saturating protein concentration; N is the Hill coefficient.

### Nucleosome NMR sample preparation and spectroscopy

The nucleosome storage buffer was exchanged with NMR buffer by dialysis. The NMR buffer was made in D_2_O and contained 70 mM NaPO_4_ (pD 7.6), 10 mM deutereted DTT. The nucleosome concentration used was 50 μM. For the experiment involving Swi6-nucleosome complex, Swi6 was buffer exchanged in the NMR buffer. Nucleosome and Swi6 were mixed in a 1:6 molar ratio to get four Swi6 molecules per nucleosome. The Swi6 concentration was saturating in the concentration regime used.

All the NMR spectra were acquired at 303K on a Bruker 600 MHz spectrometer equipped with a Cryoprobe. Recorded spectra were processed using the NMRPipe software and displayed using SPARKY^61,62^. The assignments were transferred from published spectra of *Drosophila* nucleosome by inspection^28^. The two sets of spectra were remarkably similar owing to sequence conservation between *Drosophila* and *Xenopus* histones H3 and H2B (Extended Data Fig. 4). A comparison of the chemical shift positions across *Drosophila* and *Xenopus* histone spectra suggested very small differences between the two, with the average Δδ (ppm) being <0.05 (Extended Data 4a). Buffer and temperature were selected based on optimal conditions screened by fluorescence polarization.

The concentrations of Swi6-mono-nucleosome complexes used for the NMR experiments did not display detectable aggregation or precipitation as evaluated using two different metrics. Firstly, analytical ultracentrifugation carried out at comparable concentrations of the complex did not detect higher-order species beyond four Swi6 molecules bound to a mono-nucleosome. Secondly, the NMR data did not show significant overall signal loss for up to ~60 hours of measurement time. Precipitation or aggregation is expected to result in signal loss due to overall broadening of all the cross-peak signals.

### CSD NMR

Binding experiments were carried out with 80 μM ^15^N-CSD and 160 μM H2B peptide (dissolved in H_2_O, pH adjusted to 7.8) in 150 mM KCl, 20 mM HEPES pH 7.8, 2 mM DTT. HSQC spectra were recorded at 298K on a Bruker Avance DRX500 spectrometer.

CSPs were calculated from the equation:

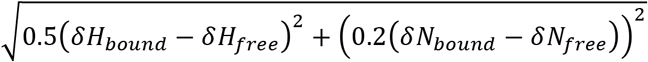

where the factor 0.2 is used as a scaling factor for nitrogen spectral width.

### HDX-MS

Solution-phase amide HDX experiments were carried out with a fully automated system described previously with slight modifications^63^. Five μl of apo-nucleosome or nucleosome-Swi6 complex (1:4) was mixed with 20 μL of D_2_O-containing HDX buffer (20 mM Hepes pH 7.5, 150 mM KCl, 10% glycerol, and 10 mM DTT) and incubated at 22°C for a range of time points (0s, 10s, 10^2^s, 10^3^s, 10^4^s or 10^5^s). Each HDX experiment was carried out in triplicate with a single preparation of each protein-ligand complex. Following on-exchange, unwanted forward or back exchange was minimized and the protein was denatured with a quench solution (5 M urea, 50 mM TCEP, and 1% v/v TFA pH 2.5) at 1:1 ratio to protein. Samples were then passed through an in-house prepared immobilized pepsin column at 50 μL min^-1^ (0.1% v/v TFA, 15 °C) and the resulting peptides were trapped on a C_8_ trap column (Hypersil Gold, Thermo Fisher). The bound peptides were then gradient-eluted (5-50% CH_3_CN w/v and 0.3% w/v formic acid) across a 1 mm × 50 mm C_18_ HPLC column (Hypersil Gold, Thermo Fisher) for 5 min at 4 °C. The eluted peptides were then analyzed directly using a high resolution Orbitrap mass spectrometer (Q Exactive, Thermo Fisher). To identify peptides, MS/MS experiments were performed on Q Exactive Orbitrap mass spectrometer over a 70 min gradient. Product ion spectra were acquired in a data-dependent mode and the five most abundant ions were selected for the product ion analysis. The MS/MS *.raw data files were converted to *.mgf files and then submitted to Mascot (Matrix Science, London, UK) for peptide identification. Peptides with a Mascot score of 20 or greater were included in the peptide set used for HDX detection. The MS/MS Mascot (Matrix Science, London) search was also performed against a decoy (reverse) sequence and false positives were ruled out. The MS/MS spectra of all the peptide ions from the Mascot search were further manually inspected and only the unique charged ions with the highest Mascot score were used in estimating the sequence coverage. Percent deuterium exchange values for peptide isotopic envelopes at each time point were calculated and processed using HDX Workbench^64^. The intensity weighted mean m/z centroid value of each peptide envelope was calculated and subsequently converted into a percentage of deuterium incorporation. This is accomplished by determining the observed averages of the undeuterated and using the conventional formula described elsewhere^65^. Corrections for back-exchange were made on the basis of an estimated 70% deuterium recovery and accounting for 80% final deuterium concentration in the sample (1:5 dilution in D_2_O HDX buffer). Deuterium uptake for each peptide is calculated for each of time points and the difference in % D values between the Apo-nucleosome and Nucleosome-Swi6 samples is shown as a heat map with a color code given at the bottom of the figure.

### Cross-linking Mass Spectrometry

Trimethyl lysine analog H3K9 mononucleosomes (85 μg, 20 μM) either with or without Swi6 (80 μM) in buffer A (20 mM Hepes, 150 mM KCl, 2 mM DTT, pH 7.5) were reacted with 25 μM EDC (added as 10x stock in buffer A) and 0.5 μM N-hydroxysulfosuccinimide (added as 15x stock in buffer A) for 60 minutes at 25°C. The reactions were quenched by adding 50 mM Tris-base and 20 mM β-mercaptoethanol and incubating for 15 minutes at 25°C. Samples were acetone precipitated and washed once with cold acetone. The pellet was resuspended in 8M Urea, 10 mM TCEP, 100 mM ammonium bicarbonate and heated at 56°C for 20 minutes, followed by alkylation with 20 mM iodoacetamide for 45 min at room temperature. The sample was diluted 6-fold with 100 mM ammonium bicarbonate and digested with 1:25 trypsin for 4 hours at 37°C followed by addition of a second aliquot of trypsin and overnight digestion.

Cross-linked peptides were desalted, fractionated by size-exclusion chromatography (SEC), and analyzed by LC-MS using a previously described method^66^. Briefly, trypsin digests were acidified to 0.2% TFA, desalted, and run over a Superdex Peptide PC 3.2/300 SEC column (GE Healthcare). SEC fractions eluting between 0.9 ml and 1.4 ml were dried and resuspended in 0.1% formic acid for LC-MS. Each fraction was separated over a 15 cm x 75 μm ID PepMap C18 column (Thermo) using a NanoAcquity UPLC system (Waters) and analyzed by a Q-Exactive Plus mass spectrometer (Thermo). The top 10 most abundant, triply charged and higher precursor ions (measured at 70,000 resolution) were selected for HCD (NCE: 24.5) and measured at 17,500 resolution.

Peak lists were generated using Proteome Discoverer 1.4 (Thermo) and searched for cross-linked peptides with Protein Prospector 5.17.2 against a target database containing Swi6 from S. *pombe* plus the four core histone sequences from *X. laevis* concatenated with a decoy database containing 10 randomized copies of each target sequence (total database size 60 sequences)^67^. Tri-methylation of histone H3 at K9 was treated as a constant modification as was loss of the initiator methionine and carbamidomethylation of cysteine. Methionine oxidation, peptide N-terminal glutamine to pyroglutamate formation, acetylation at the protein N-terminus, and mis-annotation of the monoisotopic peak (1Da neutral loss) were treated as variable modifications. EDC was designated as a heterbifunctional cross-linking reagent with specificity of aspartate, glutamate, and the protein C-terminus on one side and lysine and the protein N-terminus on the other with a bridge mass corresponding to loss of H_2_O. A mass modification range of 400-5000 Da was specified on these residues and 85 product ion peaks from the peaklist were used in the search. Precursor and product ion tolerances were 10 and 25 ppm respectively.

Cross-linked spectral matches (CSMs) were classified using a support vector machine based scoring model as described previously^66^. The final residue-pair level false discovery rate was 0.5%. Crosslink spectral counts were assigned to reside-pairs and domain-pairs in a way that normalized for ambiguous site-localization as previously detailed^66^. The dataset was then aggregated into unique cross-linked residue-pair level data with a corresponding spectral count value. Due to the prevalence of multiple, closely spaced Asp and Glu residues in a typical tryptic peptide, site-localization of EDC crosslinks is more challenging than with homobifunctional lysine-directed reagents. To address this, when the site-localization was judged to be ambiguous, all possible residue-pairs were kept with an annotation noting the ambiguity. When calculating spectral counts, fractional spectral counts were assigned to these ambiguous site localizations so that a given CSM was awarded exactly 1 spectral count. For instance, a product ion spectrum matching equally well to both K91.H4-D65.H2B or K91.H4-E68.H2B contributes 0.5 spectral counts towards each residue-pair. Decoy CSMs were retained throughout this aggregation and spectral counting process. A linear SVM model, built on five features of the Protein Prospector search output (score difference, % of product ion signals matched, precursor charge, rank of peptide 1, and rank of peptide 2) was constructed to sort cross-linked residue pairs into decoy and target classes. Crosslinked residue-pairs with an SVM score greater than 0.1, score difference greater than 5, and at least one spectral count are reported. The final residue-pair level data set is reported at specificity of 99% corresponding to 0.5% FDR.

## Data and material availability

Annotated Spectra are available using MS-Viewer at:
http://msviewer.ucsf.edu/prospector/cgi-bin/msform.cgi?form=msviewer
Using the search key: 4yyasgfaj

## Acknowledgements

We thank J. Tretyakova for help and training in histone purification; RS. Isaac for providing nucleosomal array DNA; J. Pelton and the QB3 NMR Facility at the University of California Berkeley for help with collecting and processing NMR data; M. Keenen for providing Peg-silane coated slides. We thank L. Hsieh, N. Gamarra, D. Canzio, A. Larson, E. Nora for helpful comments on the manuscript and members of the Gross and Narlikar laboratories for stimulating discussion. This work was supported by grant NIH NCATS UL1 TR000004 to SS, JDG and GJN; Sandler Family Foundation Program for Breakthrough Research Post-doctoral Fellowship to SS; NIH NIGMS R01 GM121962 to JDG; NIH R01GM108455 and GM127020 to GJN; PBBR New Frontier Research Award to GJN; Dr. Miriam and Sheldon G. Adelson Medical Research Foundation to AB, Qexactive Plus (Thermo): NIH 1S10D016229 to AB, NIH NIGMS 8P41GM103481 to AB; Instrumentation Grants-Orbitrap Fusion Lumos (Thermo): University of California, San Francisco (Program for Breakthrough Biomedical Research (PBBR).

## Author contributions

SS, JDG and GJN identified and developed the core questions. SS performed the bulk of the experiments.VD performed HDX-MS experiments and exported the data with the help of BDP. MJT performed and analyzed XLMS. SS and MJT processed HDX-MS raw data. RWT helped with the processing and collection of NMR data. SS, JDG and GJN wrote the manuscript with contributions form the other authors. GJN and JDG oversaw the project.

**Competing interests: --**

---

**Extended Data Figure 1:**
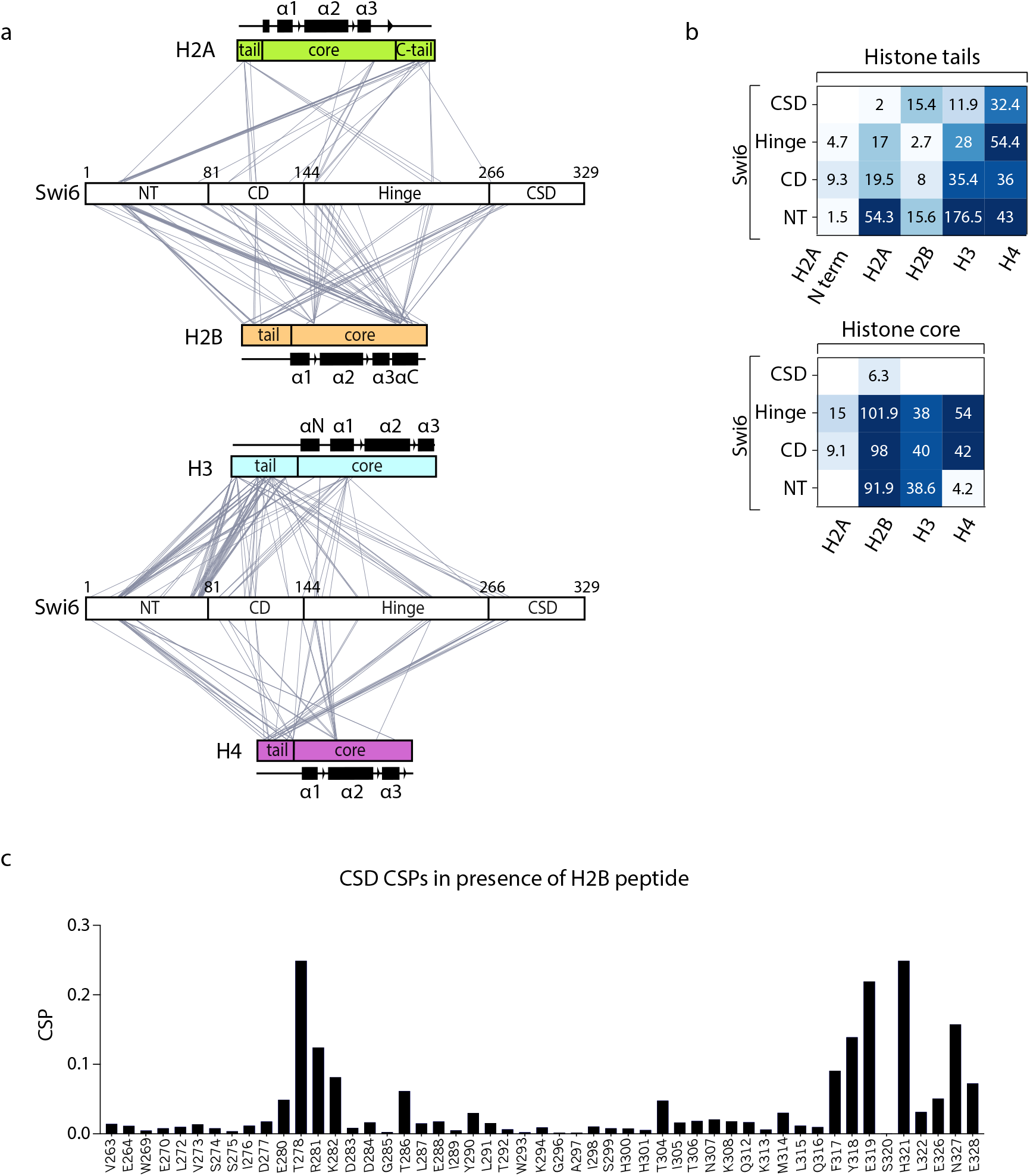
Swi6-nucleosome cross-linking analysis. (a) EDC cross-linking network between histones and Swi6. Histones tails and core regions are indicated, as well as the different Swi6 domains. (b) Interactions between Swi6 domains, histone tails, and core domains mapped by cross-link spectral counts. The number of cross-linked MS spectra matching to a given domain pair is indicated by the color of the tile as well as the numbers given. (c) Chemical shifts perturbation (CSP) for assigned resonances between Swi6 CSD alone and with the addition of the H2B peptide shown in Figure 1c.

**Extended Data Figure 2:**
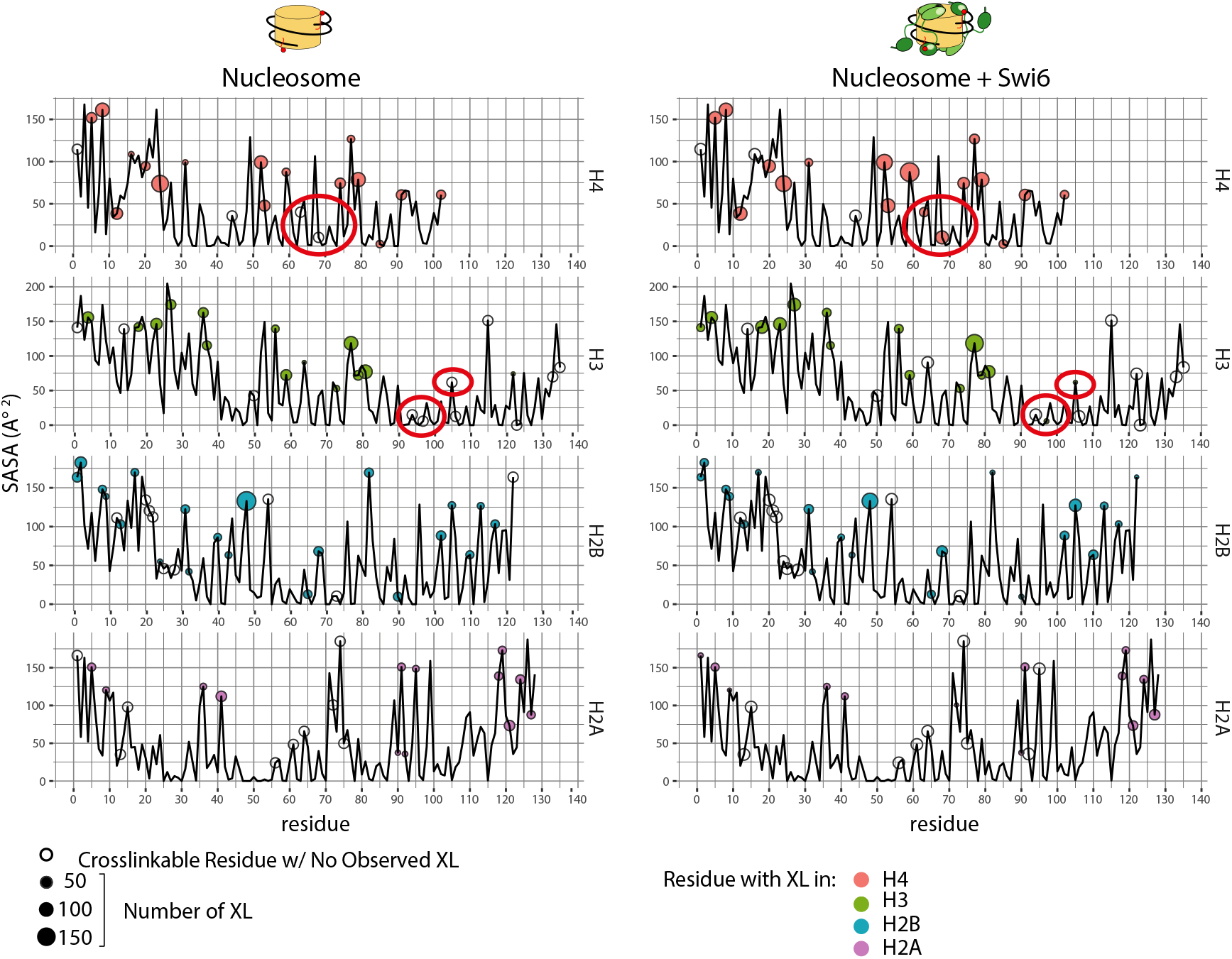
New histone-histone cross-links detected only with Swi6 binding are in buried regions of the nucleosome. The plots report the solvent-accessible surface area SASA (y-axis) vs. residues number (x-axis) for each histones. Cross-linkable residues (E, D, K) are indicated by a circle. Empty circles represent residues that do not show any cross-linking and filled circles represent cross-linked residues. The area of the circle reports the number of cross-linked spectra with one peptide mapped to a particular residue. The red ovals highlight buried residues that were observed to cross-link only in the presence of Swi6.

**Extended Data Figure 3:**
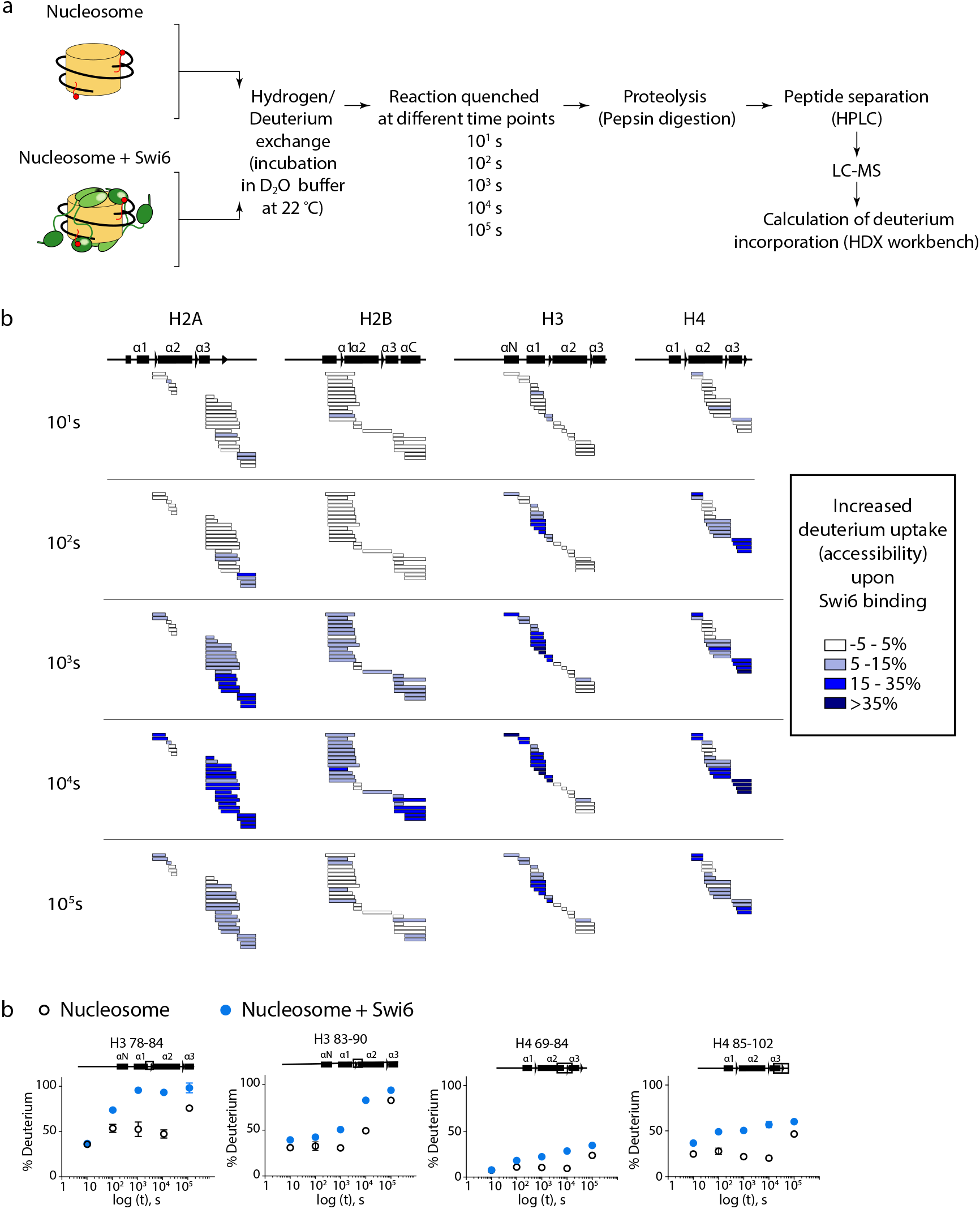
HDX-MS with nucleosome-Swi6 complex and nucleosomes alone. (a) Experimental scheme for HDX-MS. (b) Changes in deuterium incorporation is reported for every histone at five different time points. Each horizontal bar represents an individual peptide, and peptides are placed beneath schematic of secondary structure elements of the histones. Peptides are colored according to the legend, showing the mean of deprotection (n=3). (c) Kinetics of deuterium uptake of example histone peptides over the time course. Data are the mean and SD of triplicates.

**Extended Data Figure 4:**
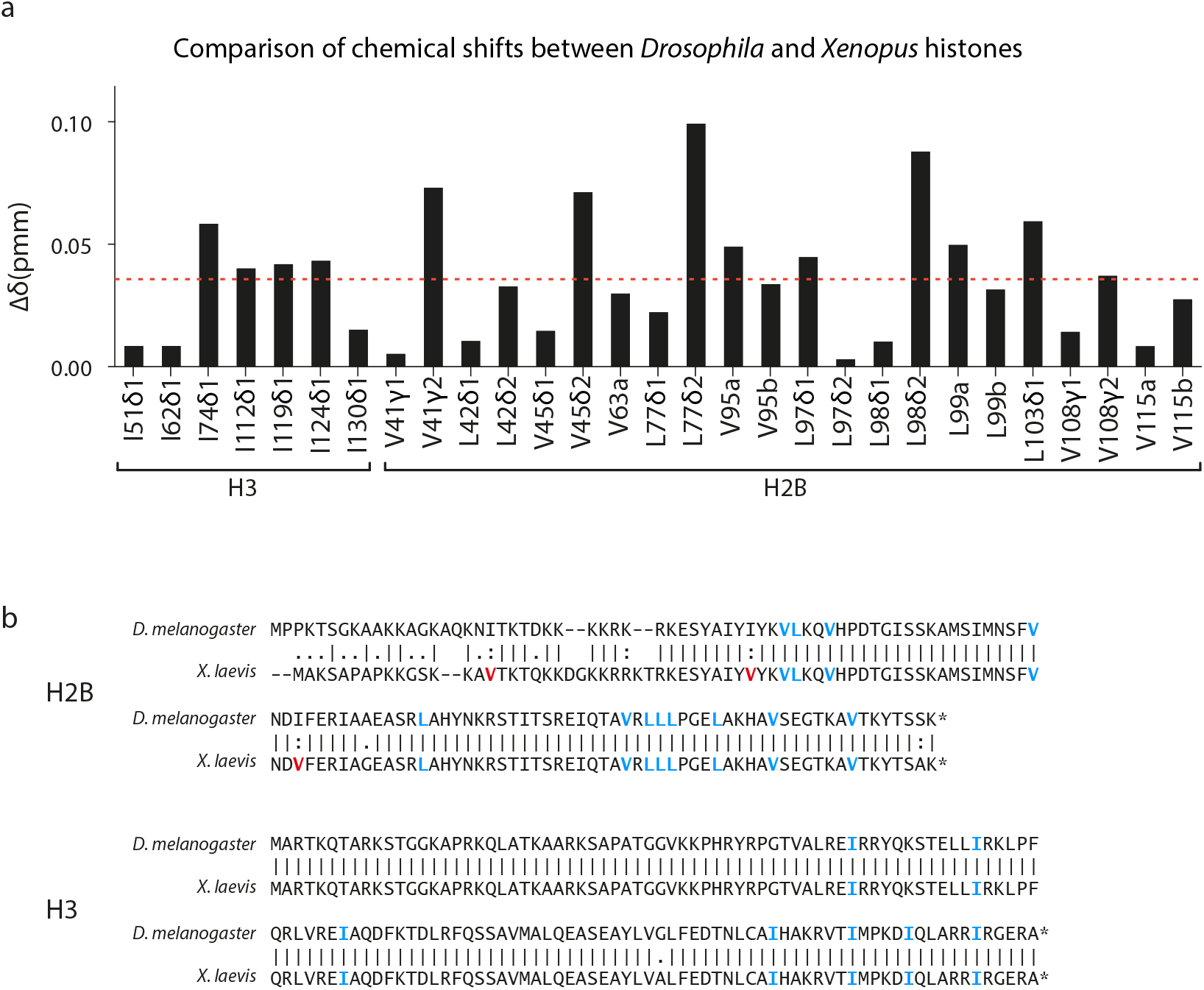
Comparison of NMR chemical shifts for ILV residues in *Drosophila* (previous work^28^) and *Xenopus* (this work) H2B and H3 histones within a nucleosome alone. (a) The Δδ plot reports the differences in chemical shift of the cross-peaks for conserved Ile, Leu and Val (ILV) residues in *Drosophila* and *Xenopus* histones. Δδ values were calculated from the equation:

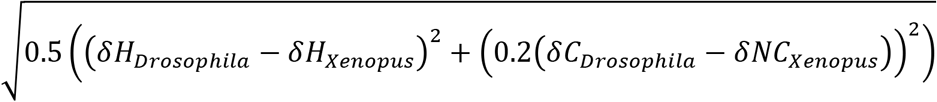 Where the factor of 0.2 is used as a scaling factor for the carbon spectral width. The average value of Δδ is shown as a red dashed line. (b) Amino acid sequence alignment of *Drosophila* and *Xenopus* H2B and H3 histones. ILV residues conserved between the two species are colored in blue, in red are the one not conserved. I residues are fully conserved between H3 histones. LV are conserved between H2B histones, except for V15, V38 and V66 that are present only in the *Xenopus* histone. V15, V38 and V66 are not included in our NMR analysis.

**Extended Data Figure 5:**
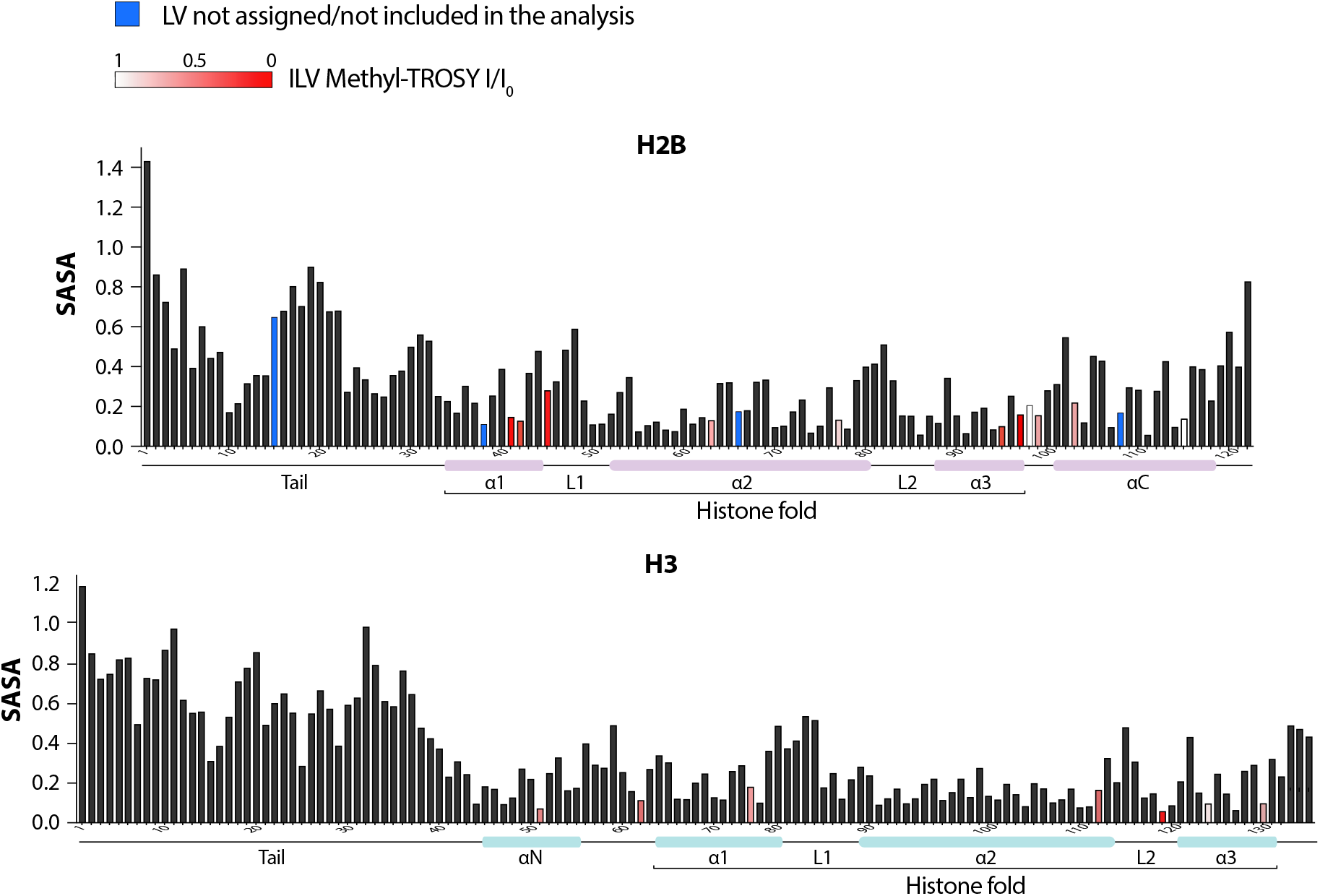
Quantification of the solvent-accessible surface area (SASA) of the residues indicated in x-axis in histone H2B (top) and H3 (bottom). SASA was calculated from the PDB structure file 1KX5 using the program POPS. The SASA values are for the entire residue and represent fraction of exposed surface area. Bars corresponding to ILV residues are colored in red with shading according to the legend, reflecting changes reported from the methyl-TROSY experiments due to Swi6 binding. ILV residues not assigned or not included in the analysis are colored in blue. The schematic of secondary structure elements is shown below the x-axis.

**Extended Data Figure 6:**
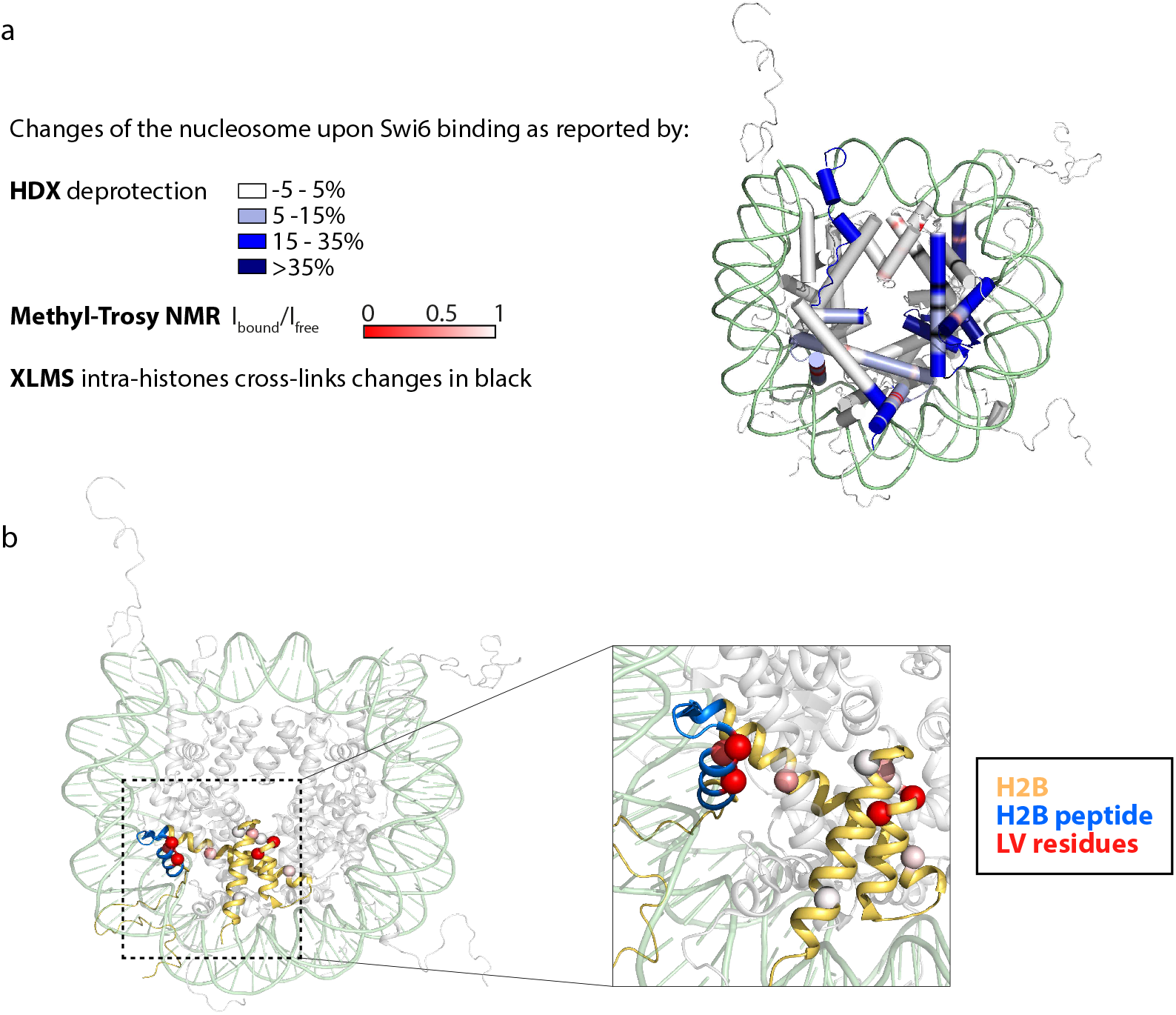
Comparison of HDX-MS, NMR and XLMS data. (a) Mapping of the Swi6-induced changes in the nucleosome structure, as detected by methyl-TROSY NMR, HDX-MS, XLMS. The figure highlights how the three different techniques consistently reported alterations in the same regions of the octamer. (b) In the nucleosome crystal structure, H2B is colored in light orange, H2B peptide used in the CSD ^1^H-^15^N HSQC NMR experiment is in cyan and Leu/Val residues analyzed in the nucleosome methyl-TROSY NMR are represented as spheres and colored according to the extent of broadening determined by Ibound /Ifree. The zoomed panel shows the location of the H2B peptide tested for binding to the CSD.

**Extended Data Figure 7:**
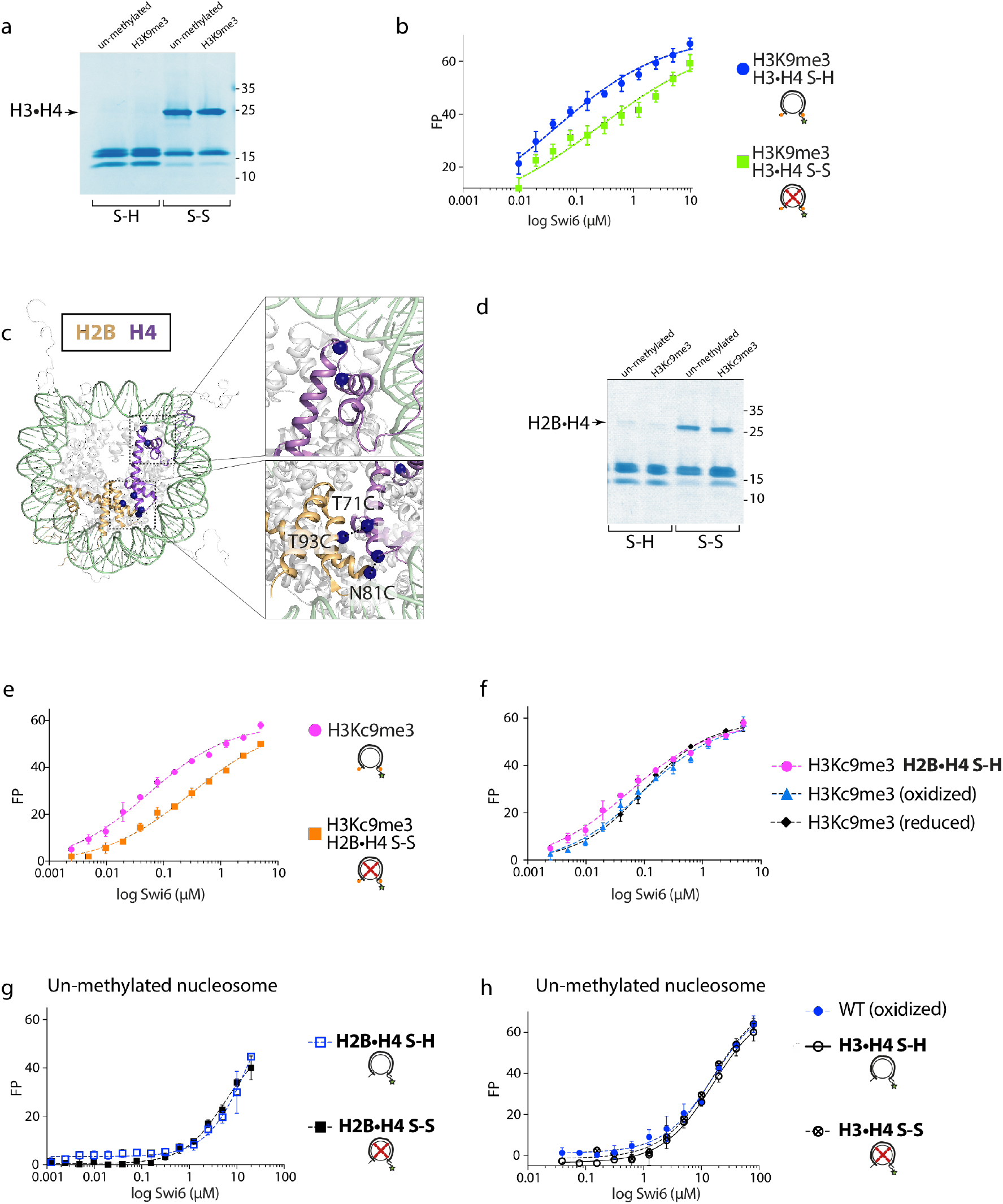
Nucleosome octamer dynamics are important for Swi6 nucleosome binding. (a) Non-reducing SDS-PAGE showing nucleosome with H3-H4 histone disulfide linked (H3•H4 S-S) or reduced (H3•H4 S-H). ~90% of H3 and H4 are disulfide-linked in both the methylated and unmethylated nucleosomes. These are the cross-linking sites used in Figure 4. (b) Nucleosome binding assays by fluorescence anisotropy showing reduced binding of Swi6 to H3K9me3 nucleosomes containing H3-H4 disulfide-linked octamer (H3•H4 S-S vs. H3•H4 S-H). Error bars represent s.e.m. (n=3). FP, fluorescence polarization units. (c) Additional residues in the H2B and H4 mutated to Cys for generating dynamically restrained octamers are represented with spheres. (d) Non-reducing SDS-PAGE showing nucleosome with H2B-H4 histone disulfide linked (H2B•H4 S-S) or reduced (H2B•H4 S-H). ~50% of H2B and H4 are disulfide-linked in both H3Kc9me3 and unmethylated nucleosomes. (e) Nucleosome binding assays by fluorescence anisotropy showing reduced binding of Swi6 to H2B-H4 disulfide-linked octamer (H2B•H4 S-S). Error bars represent s.e.m. (n=3). FP, fluorescence polarization units. (f) Fluorescence anisotropy experiments showing comparable Swi6 binding to H3cK9me3 non-oxidized, H3cK9me3 oxidized and H2B•H4 S-H mononucleosomes. These results show that the oxidation process does not alter Swi6 binding to Cys-devoid nucleosomes and that the presence of reduced Cys does not affect Swi6 binding either. (g, h) Fluorescence anisotropy measuring Swi6 binding to un-methylated nucleosome, H2B•H4 S-H and S-S (g), and H3•H4 S-H and S-S (h). Both disulfide linkages have no effect on Swi6 binding to un-methylated nucleosome. Error bars represent s.e.m. (n=3). FP, fluorescence polarization units.

**Extended Data Figure 8:**
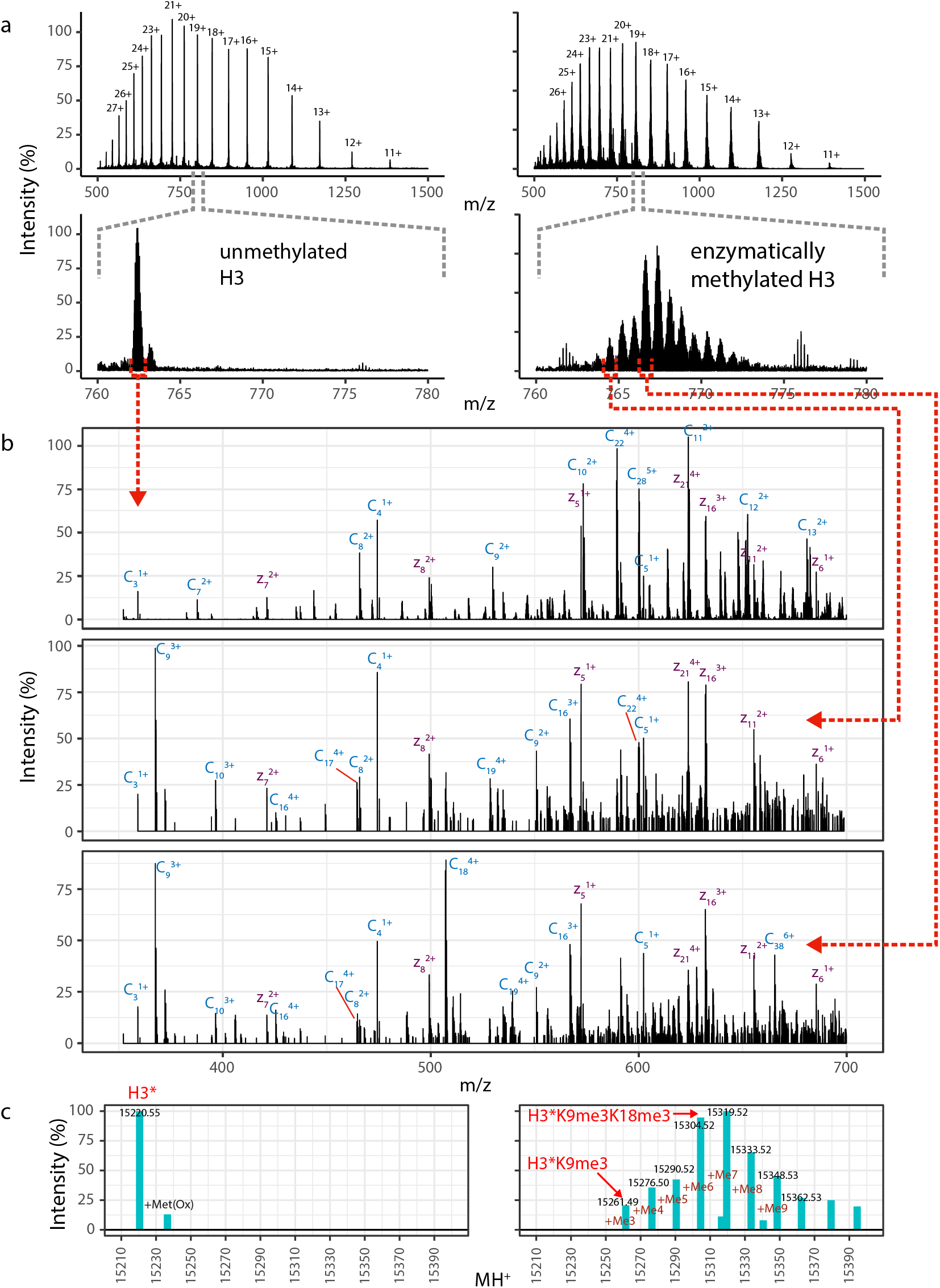
Characterization of H3 Enzymatic Methylation by Dim5. Methylation of Histone H3 was followed by online LC-MS-ETD-MS of the intact proteins. (a) Charge state envelope of untreated (left panel) and methylated (right panel) H3. The lower panel, focused on a single charge state, shows disappearance of the starting material and the formation of higher mass species, spaced 14 Da apart. (b) ETD-MS of precursor ions corresponding to un-methylated H3 (top), H3-K9me3 (middle) and H3-K9me3-K18me3 (bottom). The rationale for these assignments is based on the precursor mass values and by the product ions. Z-ions (purple) do not change between the three spectra, while the pattern of c-ions (blue) are mass shifted by 42 Da and charge-state shifted by 1+ at C9 between the top and middle/bottom panels, and again at C18 between the middle and lower panels. These assignments are further validated by bottom-up proteomics analysis of Lys-C digested samples (not shown). (c) The precursor ion spectra in (a) were deconvoluted using Xtract, which models both the charge states and the isotope distributions. Deconvoluted MH+ values are consistent with multiple methylation states (due to the difficulty of modeling isotope distributions from large proteins, particularly as there is some underlying oxidation, Xtract sometimes picks the wrong monoisotope). Deconvoluted intensities show that the enzymatically treated sample (right) contains no significant un-methylated H3 or mono- and di-methylated H3K9. While, 100% of analyzed sample is tri-methylated at H3K9, additional methylations occur at H3K18 as noted above.

**Extended Data Figure 9:**
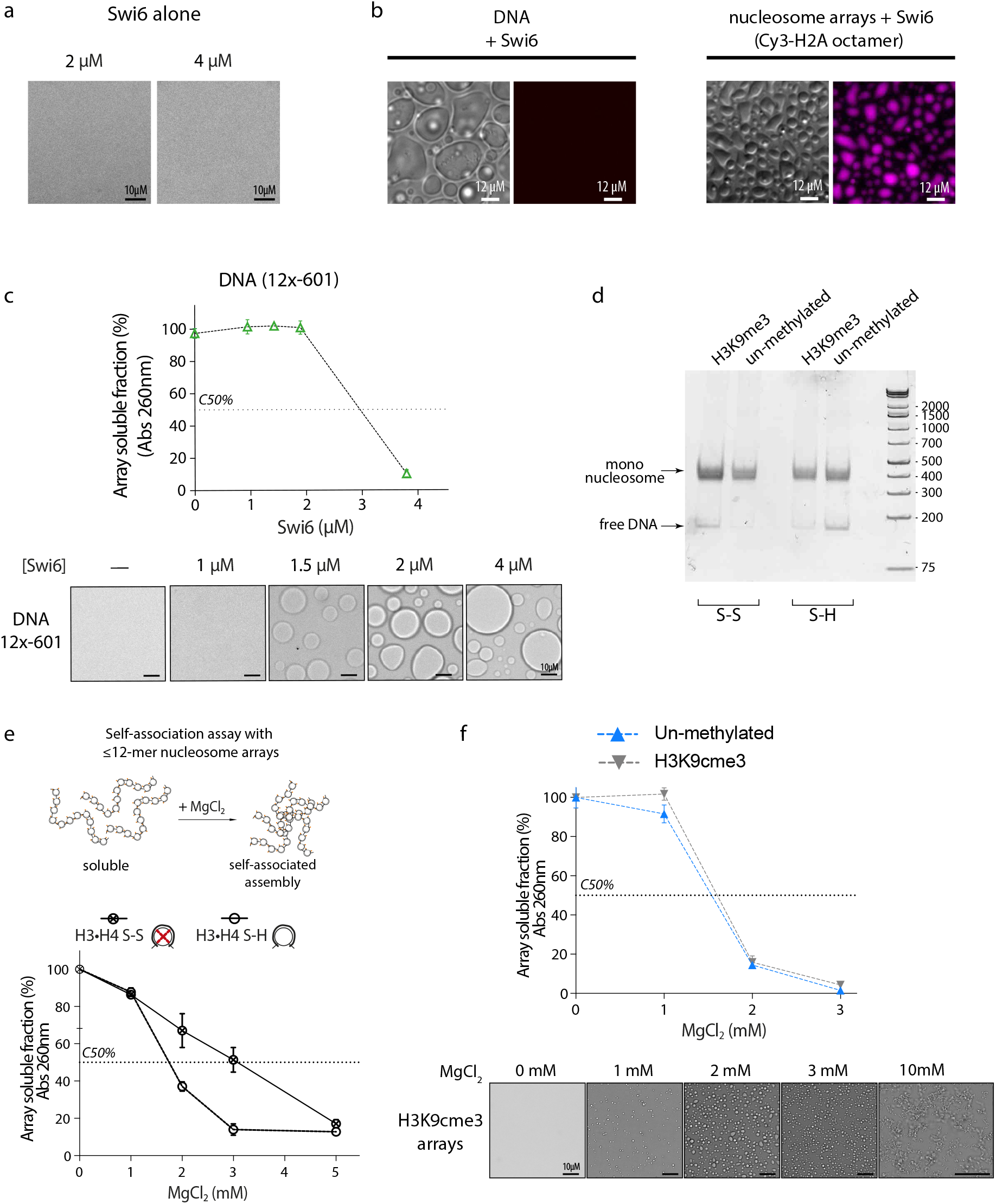
Assessment of droplet formation in different conditions. (a) Swi6 alone does not form phase-separated liquid droplets. Bright field representative images of the Swi6 alone. (b) Chromatin is included in Swi6-induced droplets. Nucleosome arrays are assembled with H2A-Cy3 and H3Kc9me3 octamers. 40 nM arrays and 4 μM Swi6. (c) Left: precipitation assay performed with naked DNA (601×12 ~2 kbp) in presence of increasing concentration of Swi6. Right: bright field representative images of the DNA-Swi6 complex analyzed in the left panel, showing the formation of phase-separated droplets. (d) Nucleosome arrays quality controls. Arrays are digested with HpaI and run on a native gel showing comparable octamer saturation and absence of over-assemblies. (e) Mg^2+^ driven precipitation of nucleosome arrays with and without disulphide cross-links between H3 and H4. (F) Top: Mg^2+^ driven precipitation of un-methylated and H3K9cme3 arrays are comparable. Bottom: Representative bright field images of the H3K9cme3 array precipitation analyzed in the upper panel, showing the formation of phase-separated droplets. The assay is performed in TE0.1 and 75mM KCl.

**Table S1.**

| Nucleosome H3K9me |  | Nucleosome |  |
| --- | --- | --- | --- |
|  | K<sub>d</sub> (μM) |  | K<sub>d</sub> (μM) |
| H3Kc9me3 | 0.18 (± 0.02) | WT | 16.43 (±1.9) |
| H3Kc9me3 **H2B•H4 S-H** | 0.12 (± 0.01) | **H2B•H4 S-H** | 13.96 (±1.8) |
| H3Kc9me3 **H2B•H4 S-S** | 0.6 (± 0.05) | **H2B•H4 S-S** | 8.8 (±0.92) |
| H3K9me **H3•H4 S-H** | 0.05 (± 0.01) | **H3•H4 S-H** | 15.8 (±1.6) |
| H3K9me **H3•H4 S-S** | 0.30 (± 0.06) | **H3•H4 S-S** | 15.46 (±1.9) |

**Table S2.**

| Nucleosome | Histone mutations |
| --- | --- |
| H3Kc9me3 | H3 C110A, K9C |
| H3Kc9me3 **H2B•H4 S-H/S-S** | H3 C110A, K9C,<br>H4 A38C, I46C, T71C, N81C, A76C, T93C |
| **H2B•H4 S-H/S-S** | H3 C110A<br>H4 A38C, I46C, T71C, N81C, A76C, T93C |
| **H3•H4 S-H** | H3 C110A, I62C<br>H4 A33C |

